# Comparative metagenomics of tropical reef fishes show conserved core gut functions across hosts and diets with diet-related functional gene enrichments

**DOI:** 10.1101/2024.05.21.595191

**Authors:** Derek G. Wu, Cassandra R. Harris, Katie M. Kalis, Malique Bowen, Jennifer F. Biddle, Ibrahim F. Farag

## Abstract

Fish gut microbial communities are important for the breakdown and energy harvesting of the host diet. Microbes within the fish gut are selected by environmental and evolutionary factors. To understand how fish gut microbial communities are shaped by diet, three tropical fish species (hawkfish, *Paracirrhites arcatus*; yellow tang, *Zebrasoma flavescens*; and triggerfish, *Rhinecanthus aculeatus*) were fed piscivorous (fish meal pellets), herbivorous (seaweed), and invertivorous (shrimp) diets, respectively. From fecal samples, a total of 43 metagenome assembled genomes (MAGs) were recovered from all fish diet treatments. Each host-diet treatment harbored distinct microbial communities based on taxonomy, with *Proteobacteria*, *Bacteroidota*, and *Firmicutes* being the most represented. Based on their metagenomes, MAGs from all three host-diet treatments demonstrated a baseline ability to degrade proteinaceous, fatty acid, and simple carbohydrate inputs and carry out central carbon metabolism, lactate and formate fermentation, acetogenesis, nitrate respiration, and B vitamin synthesis. The herbivorous yellow tang harbored more functionally diverse MAGs with some complex polysaccharide degradation specialists, while the piscivorous hawkfish’s MAGs were more specialized for the degradation of proteins. The invertivorous triggerfish’s gut MAGs lacked many carbohydrate degrading capabilities, resulting in them being more specialized and functionally uniform. Across all treatments, several MAGs were able to participate in only individual steps of the degradation of complex polysaccharides, suggestive of microbial community networks that degrade complex inputs.

**Importance:** The benefits of healthy microbiomes for vertebrate hosts include the breakdown of food into more readily usable forms and production of essential vitamins from their host’s diet. Compositions of microbial communities in the guts of fish in response to diet have been studied, but lack a comprehensive understanding of the genome-based metabolic capabilities of how they support their hosts. Therefore, we assembled genomes of several gut microbes collected from the feces of three fish species that were being fed different diets to illustrate how individual microbes can carry out specific steps in the degradation and energy utilization of various food inputs and support their host. Herbivorous fish harbored a functionally diverse microbes with plant matter degraders, while the piscivorous and invertivorous fish had microbes that were more specialized in protein degradation.

## Introduction

There are many factors that affect the composition of gut microbes in fish, including the microbial composition of the surrounding water, diet, and environmental conditions such as salinity, temperature, and pH (1, 2). Additionally, gut microbiomes are dynamic, where fish hatch with few microbes in their gut as larvae (3), which increase in quantity and diversity as they take in water and food from the surrounding environment (1). The composition of the aquatic microbial environment has been shown to correlate with the gut microbiomes of the fishes in those aquatic environments (4, 5, 6), while other studies have shown a divergence between gut and environmental microbial communities (7). Gut microbiomes in individual fishes in the same environment often possess distinct microbial communities (5), and these communities can remain relatively stable over time in the absence of disturbances (8).

In marine fish, *Proteobacteria*, *Fusobacteria*, *Firmicutes*, *Bacteroidetes*, *Actinobacteria* and *Verrucomicrobia* are among the most commonly identified phyla in intestinal samples (9). At lower taxonomic scales, *Vibrio*, *Pseudomonas*, *Achromobacter*, *Corynebacterium*, *Alteromonas*, *Flavobacterium*, and *Micrococcus* are considered to be the predominant gut colonizers (1). Although the existence of baseline similarities in core gut microbial communities among congeneric fish hosts has been suggested (10, 11, 12, 13), there are also studies that show that microbiomes between congeneric hosts can diverge based on differences in environment and trophic level (14). For example, in a study of migratory rabbitfish, less than half of the gut microbial community remained consistent across the fish’s migratory path (15). There are also reported convergences of gut microbial communities between surgeonfish and nonsurgeonfish, suggesting that core microbiomes across fish species may exist (16, 17).

Diet is an important factor that affects the gut microbiome. It has been shown that the midgut microbial community of convict surgeonfish reflects the microbial community of the algae that it was eating, suggesting a direct gut microbiome seeding related to diet (18). Yet, most diet-gut microbiome connections are more indirect. For example, larval diet supplementation provided to gilthead sea bream, European sea bass, and rainbow trout shifted their gut microbial communities in different ways, varying by host species (19). Similarly, changing the diet of sea bream from fishmeal to vegetables resulted in clear changes in microbial community composition, but not overall community diversity (20).

Gut microbial communities also vary by feeding strategy and trophic level (6, 21). In the microbial community composition of surgeonfish, herbivorous and algivorous diets are most associated with distinct communities, while carnivorous and omnivorous diets tend to have dynamic and less distinct compositions (16). Not only does diet affect the composition of microbes found in the gut, but it also affects the metabolism that occurs in the gut. Carnivorous diets are classified by the presence of pathways that relate to catabolism and biosynthesis of amino acids (22). Herbivorous diets are classified by the pathways linked to pyruvate metabolism and a high abundance of short chain fatty acids, which are indicators of fermentation happening in the gut (15, 23). Because herbivores do not intake an abundance of protein in their diet, they go through the process of fermentation to produce amino acids (22). To date, invertivorous diets have been less studied for gut microbiome functions.

Several studies have shown how individual gut microbes play putative roles in the degradation of key diet inputs (24). For example, in guts from carp fed grass diets, hundreds of bacterial isolates have been cultured that demonstrate cellulolytic activity (25). A study of three herbivorous fishes found that the activity of amylase enzymes in the guts positively correlated with diets that were higher in rhodophytes, while laminarases were negatively correlated with phaephytic diets (26). Using a genomic content estimator, PICRUSt, based on 16S rRNA amplicon libraries, and enzymatic tests, herbivorous diets were associated with fish gut microbes that produced more cellulases, while carnivorous diets were correlated with gut microbes that produced more trypsins (27). In rabbitfish, seaweed diets resulted in diverse gut microbial communities that were able to produce nonstarch polysaccharide degrading enzymes (28). However, these patterns likely vary by host species and diet, considering that there are alternative studies that have found little functional ability of microbial communities to degrade diet inputs (29).

The majority of studies on fish gut microbiomes have utilized only small subunit ribosomal gene amplicon sequencing for analysis, and they have clearly shown that microbiomes vary based on diet (14, 15, 19, 20, 30, 31). Few studies have examined in detail the metabolic pathways that exist across the gut microbiota of different fish species, which metagenomic analyses can provide unique insights into (32, 33). As such, we examined the gut inhabitants from multiple coral reef fish to probe at putative microbial functions by maintaining these wild caught fish in aquaria while controlling their diets. We examine three fish species in this study: yellow tang (*Zebrasoma flavescens*), humu humu Triggerfish (*Rhinecanthus aculeatus*), and arc-eye Hawkfish (*Paracirrhites arcatus*). Yellow tangs are herbivores that naturally feed on algae and marine plants; in the experiment they were fed seaweed, a natural herbivorous diet.

Triggerfish are invertivores that naturally feed on shrimp, clams and snails; for this experiment these fish were fed mysis shrimp. Hawkfish are generalist predators feeding on small fishes and invertebrates; they were fed fish pellets for this study. The overarching goal of this study was to determine the extent to which fish gut MAGs corresponded to selection pressures from the host diet using a comparative metagenomic approach. We hypothesized that the fish gut communities would primarily be determined by the biochemical capabilities of recovered metagenomes to degrade and harvest energy from the host’s specific diet, thus the major energy harvesting pathways exhibited by the MAGs would largely reflect the substrate inputs (i.e., fatty acid oxidation for high fat diets and mixed acid fermentation pathways for high polysaccharide diets). In this work, we show that the core potential metabolism of each fish gut microbiome is redundant across diets and species of fish, with greater accessory metabolisms for the herbivorous diet. We also show that the method by which one analyzes gut samples (i.e., 16S rRNA amplicon sequencing, metagenome construction, binning) has a strong role in determining which microbes are detected and included in downstream analyses, underscoring the importance of diverse sequence methodology for the continued study of gut microbial communities.

## Results

### Diet Composition

The majority of both the fish meal pellet (56.1%) and shrimp (69.3%) diets was composed of protein (Supplemental Table 1). The fish meal pellets had a total fat content of 16.6%, while the shrimp diet was 6.87% fat (Supplemental Table 1). Of the fats, both the fish pellets and shrimp had a broad composition of fatty acids. From the fish pellet diet, the largest contributions of fats were C22:6n3 docosahexaenoic, C20:1n9 cis eicosenoic, and C16:1n7 palmitoleic acids (Supplemental Table 2). From the shrimp diet, the largest contributions of fats were C16:0 palmitic, C22:6n3 docosahexaenoic, and C20:5n3 eicosapentaenoic acids (Supplemental Table 2). Both diets had a small component of fiber (2.42% and 3.57% for fish pellets and shrimp, respectively) (Supplemental Table 1). Both diets could provide gut microbes with biologically relevant minerals, such as potassium, calcium, phosphorus, and metals like iron, magnesium, zinc, manganese, and copper (Supplemental Table 1). The seaweed, fed as nori paper, contained almost equivalent protein and carbohydrates (5.7% and 5%, respectively), and minor components of fiber and sugar (both 0.4%). Seaweed had large amounts of iodine, vitamins C, A and B12 and also contained calcium, potassium, iron and magnesium.

### Initial 16S rRNA gene amplicon and metagenome communities

We first examined the fish comparatively using 16S rRNA gene amplicon sequencing, which identifies only the taxonomic representation of the sample. With this analysis, the hawkfish gut community was least diverse, and all samples showed an abundance of *Enterobacteriales* within the *Gammaproteobacteria* (Supplemental Figure 1). We then proceeded with metagenomic analysis of a later timed sample. Using coverage of a single copy ribosomal gene, *rps3*, as our taxonomic and abundance indicator, we saw that the microbial composition varies, with a high representation of *Vibrionales* in each sample that was not initially observed by amplicons (Figure 1). The greatest diversity of taxa was within the tang sample, which was also not observed in amplicon data (Figure 1).

**Figure 1:**
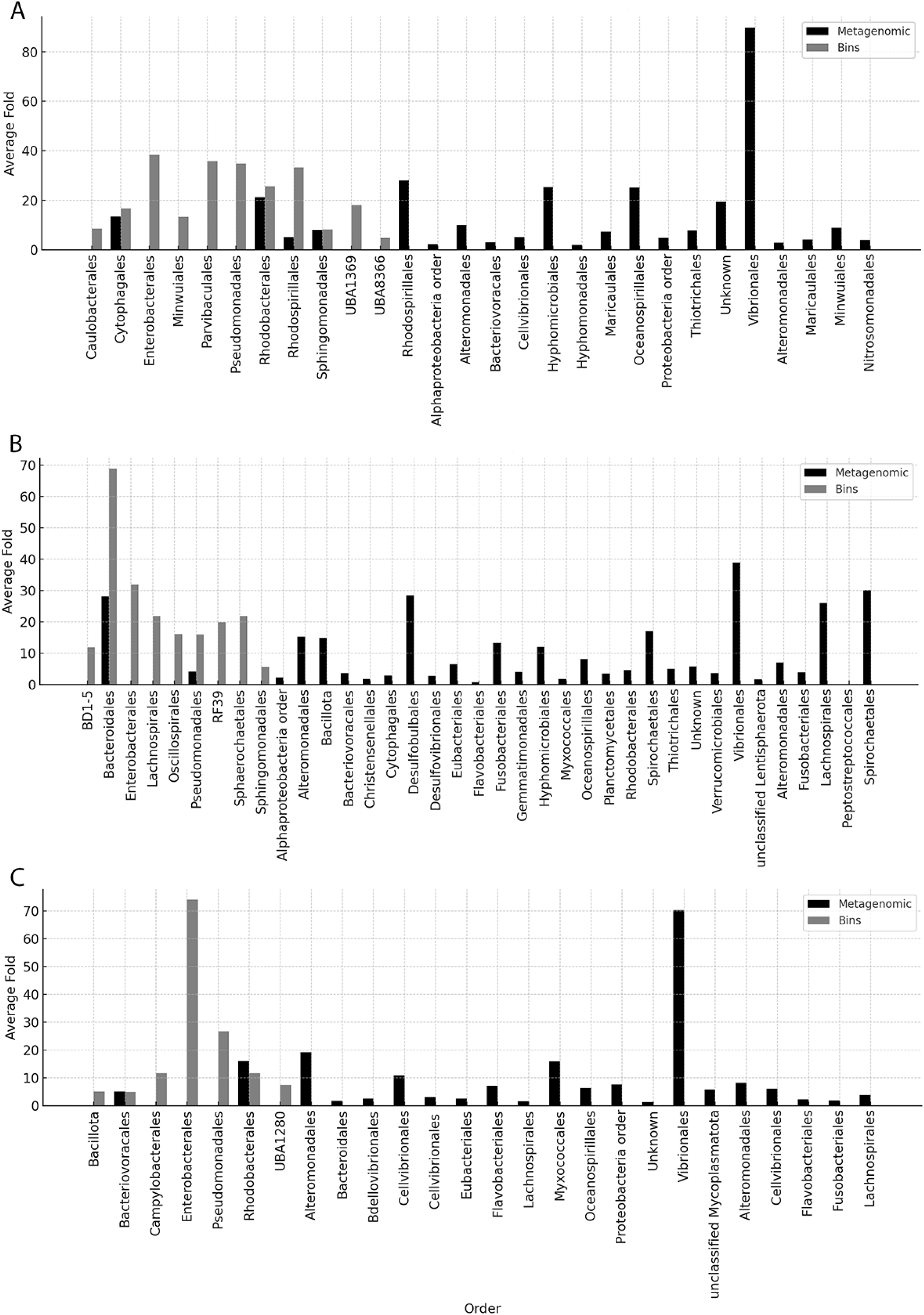
Coverage of 30S ribosomal protein annotations from the assembled metagenomes compared to coverage within each MAG. Samples shown are a) piscivorous hawkfish, b) herbivorous yellow tang, and c) invertivorous triggerfish. The comparison highlights which taxa are represented by MAGs. Presumably those not in MAGs have a high degree of species diversity and could not be accurately binned. The taxonomy of the 30S ribosomal protein gene was determined by best hit in Uniprot, the taxonomy of the MAG was determined by GTDB.

### Metagenome-Assembled Genome (MAG) Recovery

After assembly and binning (Table 1), we recovered MAGs, quality checked and identified them (Table 2). A total of 38 MAGs were recovered from the piscivorous hawkfish treatment, and 18 passed our quality threshold with an average genome completeness and contamination of 91.44% and 4.47%, respectively (Table 2). These 18 MAGs contained 4 taxonomic classes: *Alphaproteobacteria* (8), *Bacteroidia* (3), *Gammaproteobacteria* (6), and *Gracilibacteria* (1) (Table 2). The MAGs recovered were, for the most part, not reflective of the potential diversity of each sample (Figure 1), as the Vibrionales in each sample were likely too diverse to allow the MAG clustering. However, *Bacteroidales* were recovered in the tang sample and were also abundant in the unassembled metagenome (Figure 1) and the amplicon data (Supplementary Figure 1). *Enterobacterales*, which was abundant in the amplicon data and MAG data for the triggerfish, were not seen in the unassembled metagenome (Supplementary Figure 1; Figure 1). These comparisons show how the microbial community can seem to shift based on the analysis method used and highlights how studies of fish guts are best done using multiple methods of analysis. Since our MAG analysis is not comprehensive of the total microbial community, we take each analysis as a first-pass in the comprehensive metabolic analysis of fish guts.

**Table 1:** Metagenome statistics.

|  | Dataset | Size (bp) | Contigs in assembly | Max contig (bp) | Ave contig (bp) | N50 (bp) | Bins generated<sup>1</sup> |
| --- | --- | --- | --- | --- | --- | --- | --- |
| Hawkfish | H1N1 | 371954018 | 791521 | 31299 | 470 | 459 | 18 |
| Tang | T4LTN | 332236806 | 687584 | 125302 | 483 | 474 | 18 |
| Triggerfish | Tr3LTN | 310457717 | 640083 | 67851 | 485 | 471 | 7 |

<sup>1</sup>Bins that passed quality threshold

**Table 2:**
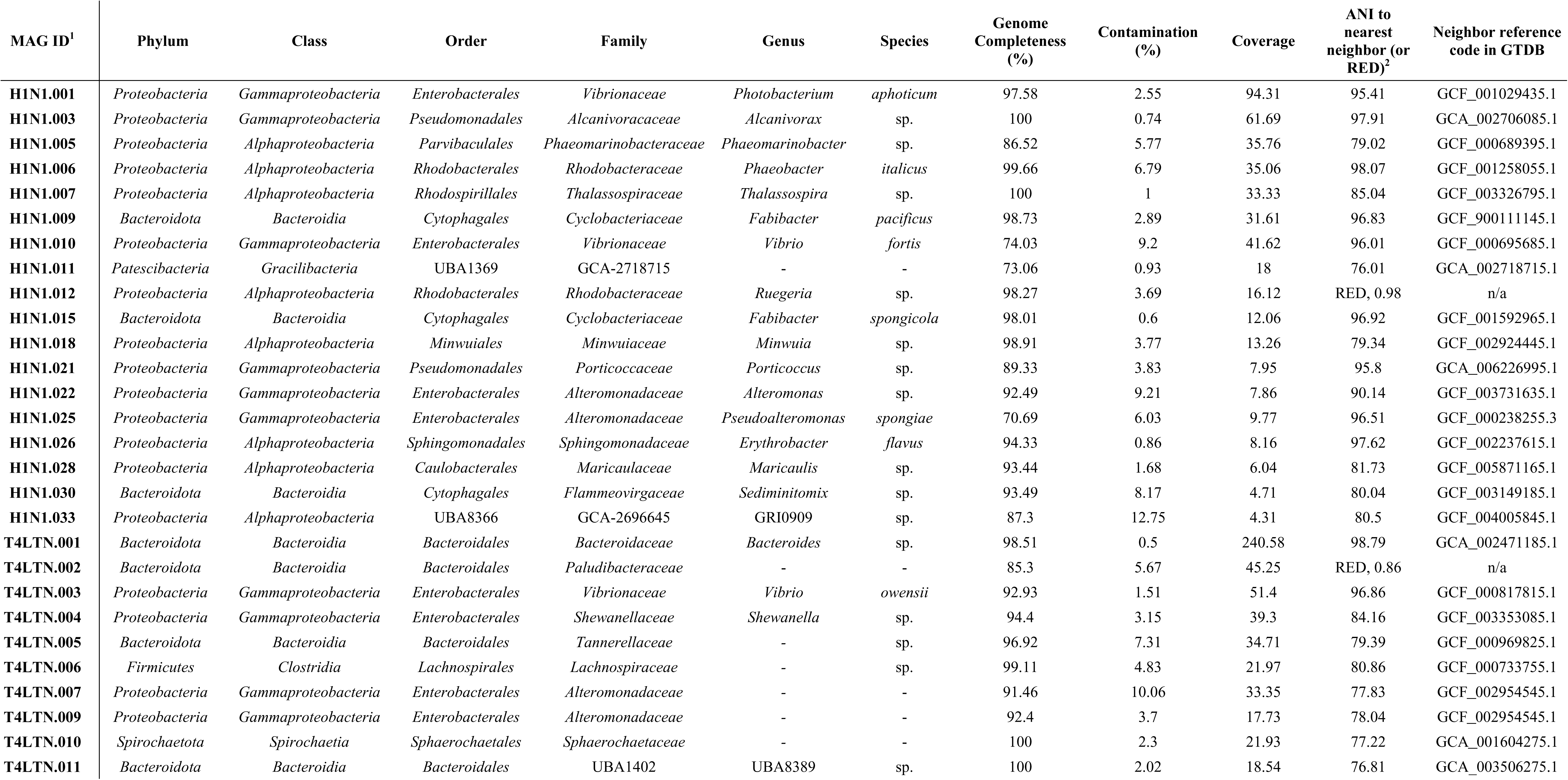
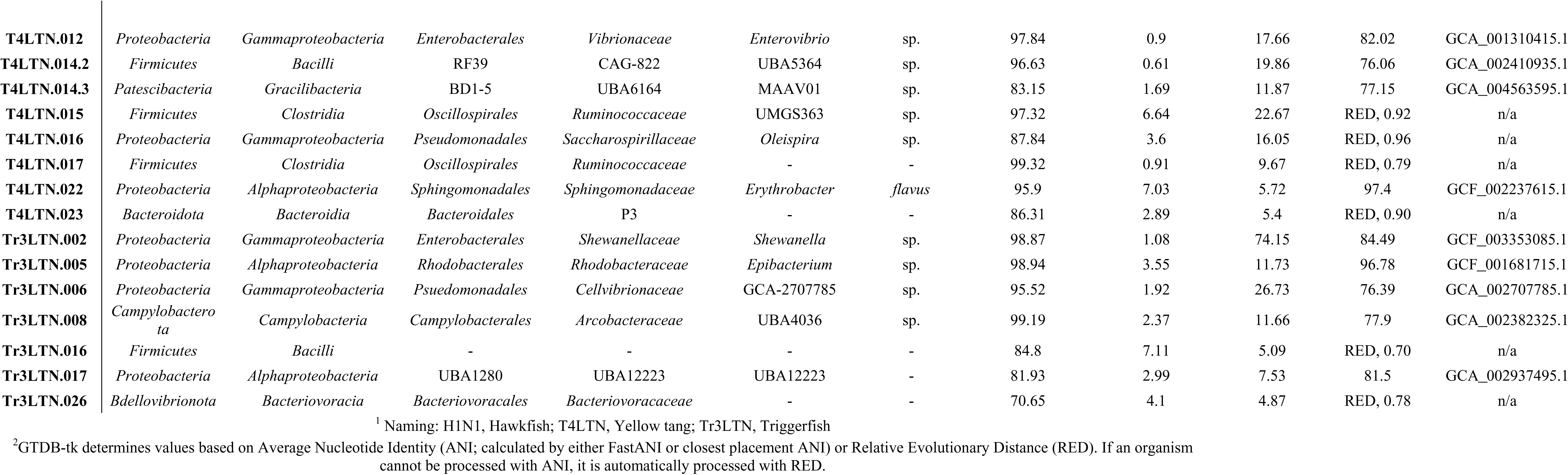
MAG taxonomic assignments, completeness, contamination, coverage and ANI values.

Our MAG analysis does highlight the ability to see microbial groups that may be overlooked due to primer bias in 16S rRNA amplicon studies or in a representation that allows them to be overlooked in an assembled metagenome. Additionally, we are able to determine how similar these MAG genomes are to known genomes. Of these MAGs, 52% are within the similarity metrics to be considered the same as previously discovered species (average nucleotide identity; ANI >95), leaving the remainder to be potentially new species or genera (Table 2). In particular, the *Gracilibacteria* MAG is within the *Patescibacteria* phylum at only 76 ANI to its neighbors. While this phylum has been seen in numerous fish guts, this MAG can only be classified to the family level.

From the herbivorous yellow tang treatment, 18/62 MAGs were also recovered and passed quality checks, with an average genome completeness and contamination of 94.19% and 3.63%, respectively (Table 2). In contrast to the piscivorous gut community, the herbivorous community appeared to have more taxonomic diversity at higher taxonomic ranks (e.g., phylum or order levels). The 18 MAGs represented 7 taxonomic classes: *Alphaproteobacteria* (1), *Bacilli* (1), *Bacteroidia* (5), *Clostridia* (3), *Gammaproteobacteria* (6), *Gracilibacteria* (1), and *Spirochaetia* (1) (Table 2). The phylogenetic novelty of this sample was also higher, with only 23% of MAGs being classified within a known species, and the majority of MAGs were only able to be classified to the family level. Some of the most novel MAGs are within the *Patescibacteria, Firmicutes* and *Bacteroidota,* based on MAG ANI values (Table 2).

Despite a similar sized dataset and what appears to be a similar assembly (Table 1), only 7/33 MAGs passed our quality threshold and were recovered from the invertivorous triggerfish treatment, with an average genome completeness and contamination of 90.0% and 3.3%, respectively (Table 2). The 7 MAGs represented 5 taxonomic classes: *Alphaproteobacteria* (2), *Bacilli* (1), *Bacteriovoracia* (1), *Campylobacter* (1), and *Gammaproteobacteria* (2) (Table 2). Only one of these MAGs was within a known species, with the majority of MAGs only being classified to the family level and one classified only to class (MAG Tr3LTN.016; *Bacillus;* Table 2). The taxonomic novelty of the MAGs within this project highlights the need for genome-level understanding of fish gut communities, beyond the small subunit ribosomal gene. Further description of novel MAGs was not done for this study, as we focused on function across diet types.

### Diet-Based Analyses

With the MAGs recovered, we analyzed each sample and its genomes for a response to diet types by performing principal component analysis on the copy numbers of enzymes within each MAG, normalizing by genome completeness. These data suggest that fish host diet may be a major driving selection factor for MAG composition (PERMANOVA; F = 6.3078; p = 0.003) (Figure 2). Hemicellulose, cellulose, chitin, and starch were forces loaded towards the herbivorous diet whereas fatty acids and proteins were loaded towards the piscivorous and invertivorous diets (Figure 2). The herbivorous gut community displayed a greater variance in gene copies, while the invertivore gut community had the narrowest variation in substrate degradation gene compositions (homogeneity of dispersion; F = 9.3049; p = 0.001) (Figure 2). Although several of the MAGs collected from the herbivorous yellow tang demonstrated specialization for the degradation of complex polysaccharides, it also contained MAGs that were more specialized in protein and fatty acid degradation. In contrast, the invertivorous triggerfish MAGs had a narrower range of substrate degradation functions, suggestive of a more specialized community that is adapted to the high protein shrimp diet.

**Figure 2:**
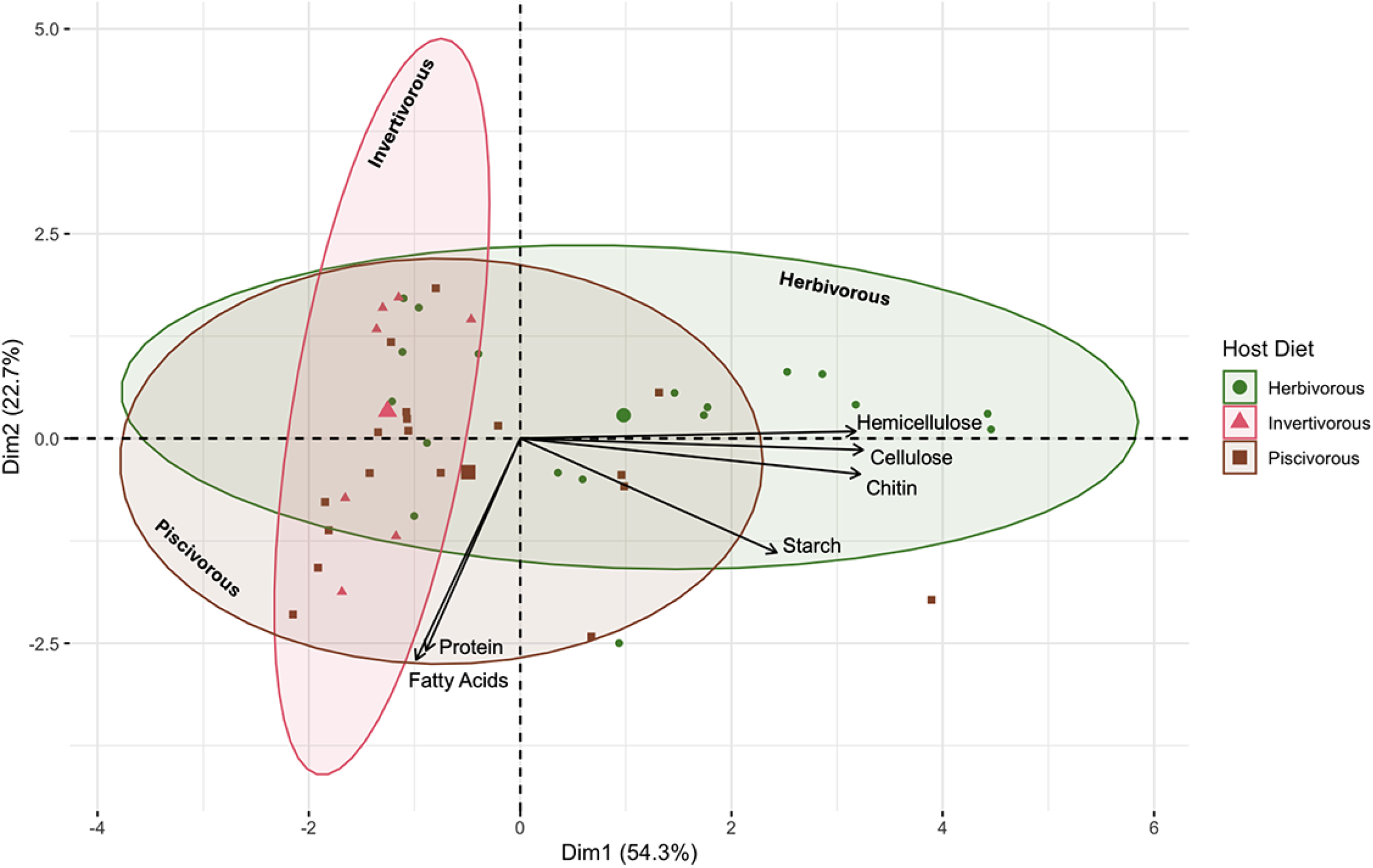
Principal component analysis of normalized enzyme gene copy numbers across all MAGs recovered from the three diet treatments. The total numbers of enzymes of each degradation pathway (proteins, cellulose, hemicellulose, starch, and chitin) normalized to the MAG’s genome completeness were used to generate components. Fatty acids were included as a binary where 1 designated that the MAG could complete all four steps of beta oxidation, while 0 designated that it could not. Ellipses represent t distributions around the centroids. Ellipses are labeled for clarity. Individual points represent individual MAGs. The larger shapes in the ellipses are group centroids.

### Carbohydrates

Overall genes encoding for carbohydrate degrading enzymes are more enriched in the MAGs recovered from herbivorous fish compared to fishes following other diets. For example, the degradation of cellulose could be completed in both the herbivorous and piscivorous diets (Figure 3a). However, MAGs recovered from the herbivorous yellow tang had significantly more gene copies for enzymes involved in the cellulose degradation pathway than both piscivorous and invertivorous hosts (Kruskal Wallis, p = 0.01) (Figure 4a). Within the piscivorous hawkfish gut, 7/18 MAGs [005, 009, 011, 022, 028, 030, and 033] could facilitate the hydrolysis of cellulose to glucose monomers (Figure 3a). Of these 7 MAGs, 4 are *Alphaproteobacteria,* 2 are *Bacteroidia*, and 1 is a *Gammaproteobacteria*. From the herbivorous diet, 9/18 MAGs [001, 002, 005, 006, 007, 009, 014.3, 015, and 016] could catalyze these reactions. Of these 9 MAGs, 3 are *Gammaproteobacteria*, 3 are *Bacteroidia*, 2 are *Clostridia*, and 1 is *Gracilibacteria*. In contrast, no MAGs recovered from the invertivorous triggerfish could degrade cellulose to simple monomers (Figure 3a). Additionally, in all treatments, there were several MAGs that could potentially catalyze at least one step of the larger cellulose degradation pathway but not the entire process, hereby referred to as “partial degraders” (Figure 3a). From the piscivorous treatment, MAGs 001, 010, and 015 were partial degraders, while from the herbivorous treatment, MAGs 003, 011, 012, and 017 were partial degraders of cellulose. The existence of these MAGs suggests that they may be opportunists that require other MAGs to initiate the breakdown of cellulose into simpler substrates for them to use.

**Figure 3:**
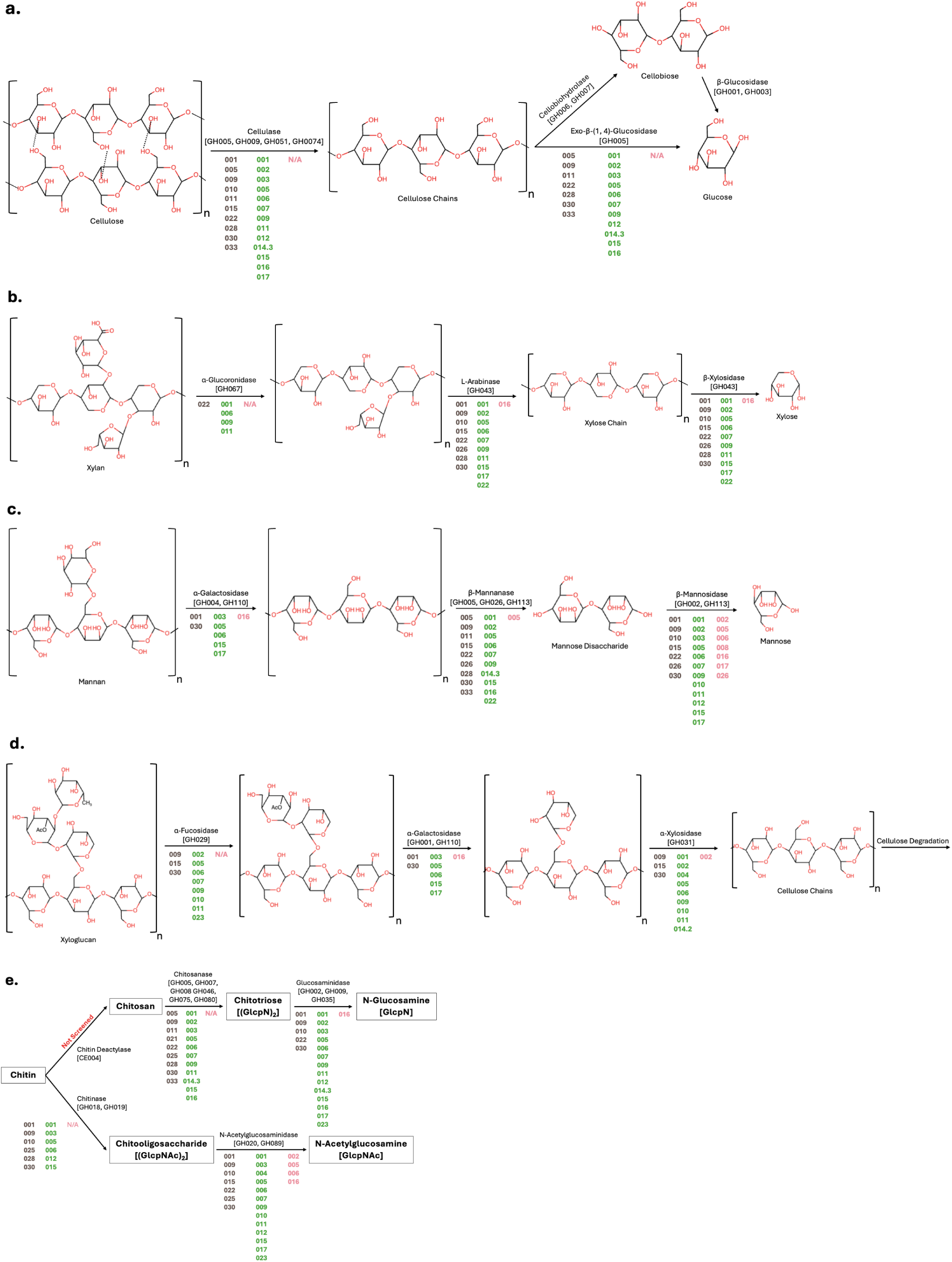
Polysaccharide degradation pathway to monosaccharide products. Shown are a) cellulose, b) xylan, c) mannan, d) xyloglucan, and e) chitin. Enzymes and their CAZyme IDs are noted above arrows connecting substrates and products. MAGs that possess genes for each enzyme are denoted beneath the arrow, where dark brown (left), green (center), and pink (right) font represent the piscivorous, herbivorous, and invertivorous treatments, respectively.

**Figure 4:**
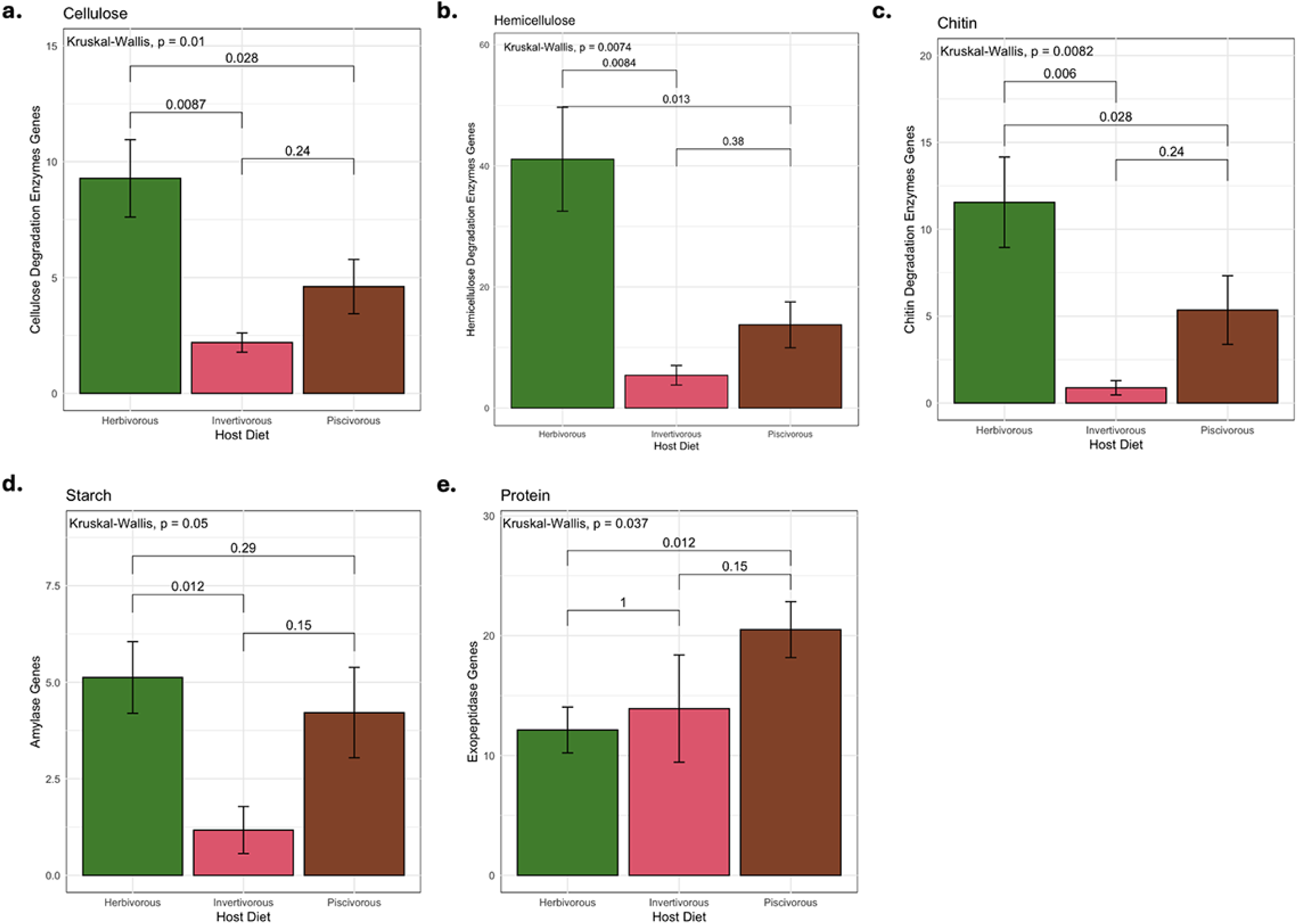
Number of substrate degradation genes per MAG normalized to genome completeness. Shown are a) cellulose, b) hemicellulose, c) chitin, d) starch, and e) proteins in each diet treatment. Kruskal Wallis values above bars represent group comparisons. Pairwise comparison values denoted by brackets are the result of post-hoc Wilcoxon tests between diets.

The MAGs recovered from the herbivorous yellow tang had significantly higher gene copies of enzymes involved in hemicellulosic degradation than both the piscivorous and invertivorous treatments (Kruskal Wallis, p = 0.0074) (Figure 4b). From the herbivorous treatment, 6/18 MAGs [001, 005, 006, 007, 011, and 015] could completely degrade at least one of the three screened hemicelluloses (Figure 3b-d). Of these 6 MAGs, 2 are *Clostridia*, 3 are *Bacteroidia*, and 1 is a *Gammaproteobacteria*. From the piscivorous treatment, only 2/18 [022 and 030] were able to fully degrade at least one hemicellulosic compound (Figure 3b-d). No MAGs recovered from the invertivorous treatment could completely degrade hemicellulose (Figure 3b-d). However, in all treatments, there were several MAGs that were partial degraders of hemicellulose backbone or one of the side chains (Figure 3b-d). From the herbivorous treatment, all MAGs except the complete degraders could be classified as partial degraders of various hemicelluloses, wherein they were capable of performing one or more steps in a hemicellulose degradation pathway. Similarly, all the MAGs recovered from the piscivorous diet could catalyze at least one step of a hemicellulose degradation pathway; however, there was a lack of capacity to hydrolyze and ferment many of the auxiliary sugars from the hemicellulose compounds.

The herbivorous MAGs had significantly greater numbers of genes involved in the degradation of chitin than both the invertivorous and piscivorous samples: 2.2 fold and 13.1 fold, respectively (Kruskal Wallis, p = 0.0082) (Figure 4c). In fact, the invertivorous MAGs showed an inability to catalyze the initial steps to either of the chitin degradation pathways (Figure 3e). Meanwhile, 2/18 MAGs [009 and 030] from the piscivorous treatment and 5/18 MAGs [001, 003, 005, 006, and 015] from the herbivorous treatment could fully catalyze the breakdown of chitin to simpler glucosamine units. The 2 MAGs from the piscivorous treatment that can degrade chitin are both in the *Cytophagales* order within the *Bacteroidia* class. Of the 5 chitin-degrading MAGs from the herbivorous treatment, 2 are *Bacteroidia*, 2 are *Clostridia*, and 1 is a *Gammaproteobacteria*.

All three samples had MAGs that encoded amylases for the degradation of starches. However, the herbivorous gut sample had significantly more amylase gene copies per MAG than the piscivorous sample (Kruskal Wallis, p = 0.05) (Figure 4d).

### Proteins

All three hosts contained MAGs that produced exopeptidases for protein degradation, allowing for basal levels of protein breakdown in the gut environment. The piscivorous gut MAGs had 1.69 fold more exopeptidase gene copies per MAG than those of the herbivorous gut community (Figure 4e), suggesting that protein degradation pathways were enriched in response to the high protein fish pellet diet. From the piscivorous community, 16/18 MAGs had more than 10 exopeptidase genes, while 11/16 and 3/7 MAGs from the herbivorous and invertivorous guts had more than 10 exopeptidase genes, respectively (Table 3). Additionally, MAG 005 from the piscivorous gut (*Phaeomarinobacter* sp.) was the only MAG recovered across the fish treatments to have a gene for collagenase, which could highlight a specialized niche for collagen degradation in response to the fish pellet diet.

**Table 3:**
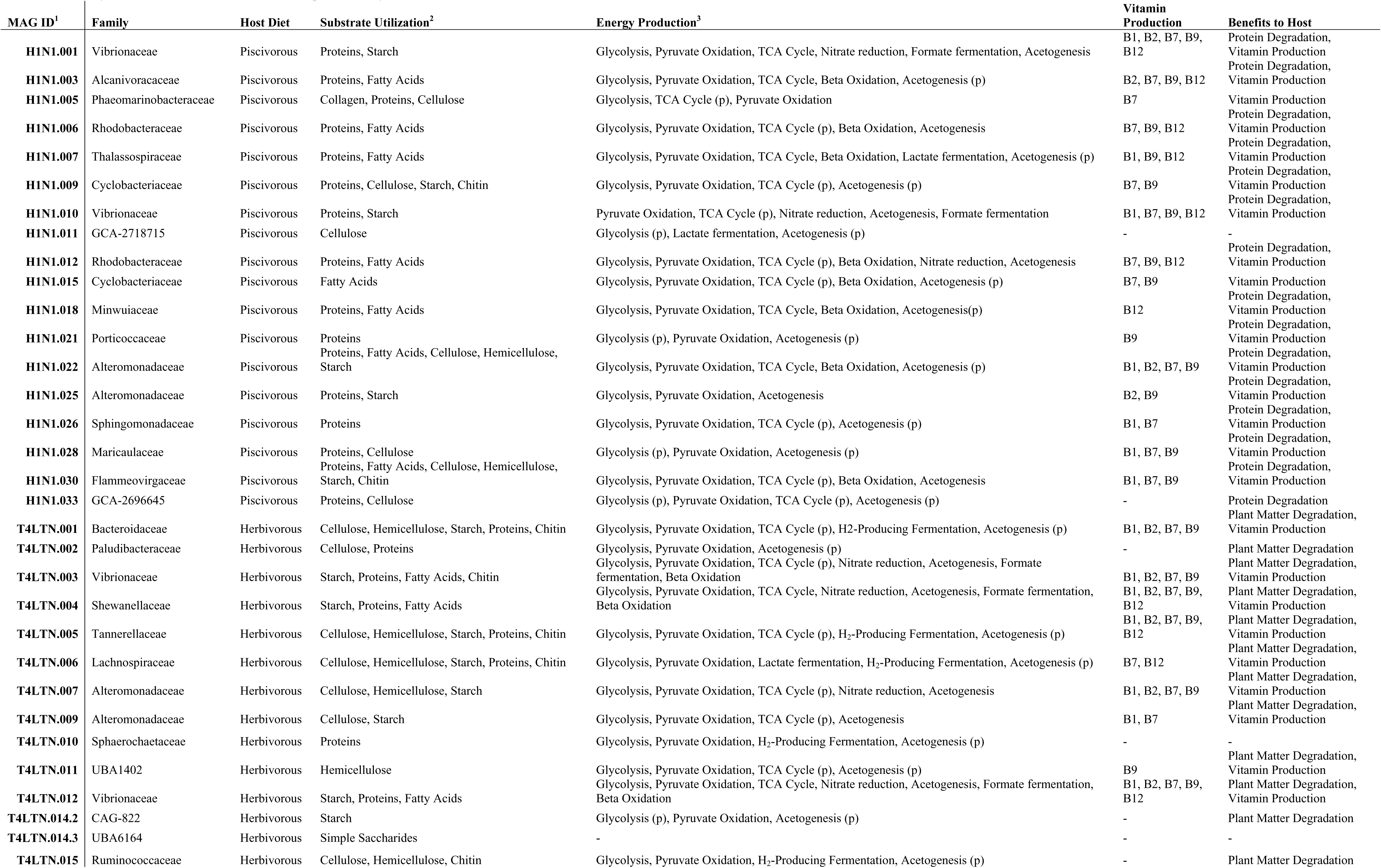
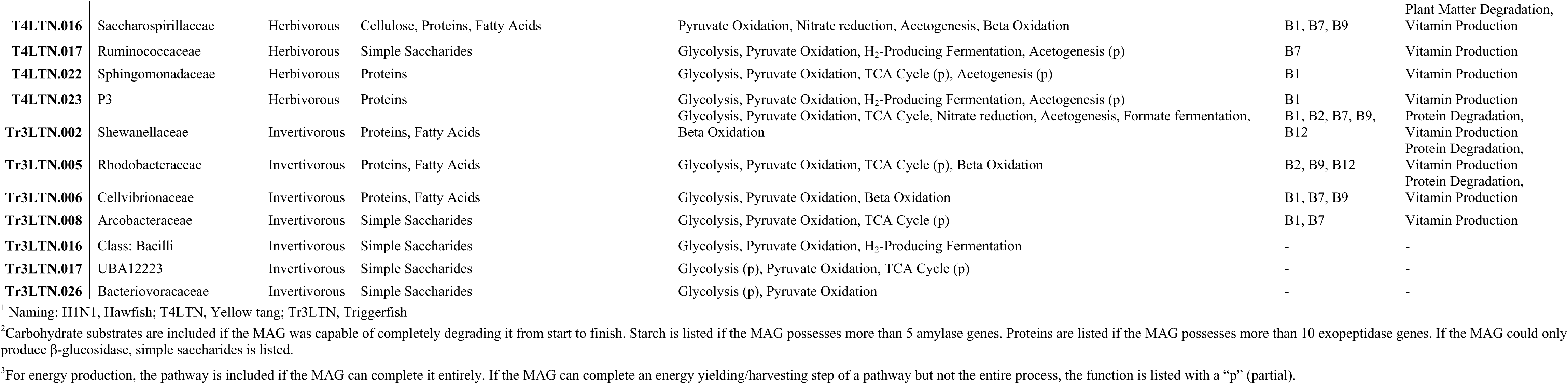
Summary of biochemical functions that could be performed by all recovered MAGs.

| MAG ID<sup>1</sup> | Family | Host Diet | Substrate Utilization<sup>2</sup> | Energy Production<sup>3</sup> | Vitamin Production | Benefits to Host |
| --- | --- | --- | --- | --- | --- | --- |
| H1N1.001 | Vibrionaceae | Piscivorous | Proteins, Starch | Glycolysis, Pyruvate Oxidation, TCA Cycle, Nitrate reduction, Formate fermentation, Acetogenesis | B1, B2, B7, B9, B12 | Protein Degradation, Vitamin Production |
| H1N1.003 | Alcanivoracaceae | Piscivorous | Proteins, Fatty Acids | Glycolysis, Pyruvate Oxidation, TCA Cycle, Beta Oxidation, Acetogenesis (p) | B2, B7, B9, B12 | Protein Degradation, Vitamin Production |
| H1N1.005 | Phaeomarinobacteraceae | Piscivorous | Collagen, Proteins, Cellulose | Glycolysis, TCA Cycle (p), Pyruvate Oxidation | B7 | Vitamin Production |
| H1N1.006 | Rhodobacteraceae | Piscivorous | Proteins, Fatty Acids | Glycolysis, Pyruvate Oxidation, TCA Cycle (p), Beta Oxidation, Acetogenesis | B7, B9, B12 | Protein Degradation, Vitamin Production |
| H1N1.007 | Thalassospiraceae | Piscivorous | Proteins, Fatty Acids | Glycolysis, Pyruvate Oxidation, TCA Cycle, Beta Oxidation, Lactate fermentation, Acetogenesis (p) | B1, B9, B12 | Protein Degradation, Vitamin Production |
| H1N1.009 | Cyclobacteriaceae | Piscivorous | Proteins, Cellulose, Starch, Chitin | Glycolysis, Pyruvate Oxidation, TCA Cycle (p), Acetogenesis (p) | B7, B9 | Protein Degradation, Vitamin Production |
| H1N1.010 | Vibrionaceae | Piscivorous | Proteins, Starch | Pyruvate Oxidation, TCA Cycle (p), Nitrate reduction, Acetogenesis, Formate fermentation | B1, B7, B9, B12 | Protein Degradation, Vitamin Production |
| H1N1.011 | GCA-2718715 | Piscivorous | Cellulose | Glycolysis (p), Lactate fermentation, Acetogenesis (p) | - | - |
| H1N1.012 | Rhodobacteraceae | Piscivorous | Proteins, Fatty Acids | Glycolysis, Pyruvate Oxidation, TCA Cycle (p), Beta Oxidation, Nitrate reduction, Acetogenesis | B7, B9, B12 | Protein Degradation, Vitamin Production |
| H1N1.015 | Cyclobacteriaceae | Piscivorous | Fatty Acids | Glycolysis, Pyruvate Oxidation, TCA Cycle (p), Beta Oxidation, Acetogenesis (p) | B7, B9 | Vitamin Production |
| H1N1.018 | Minwuiaceae | Piscivorous | Proteins, Fatty Acids | Glycolysis, Pyruvate Oxidation, TCA Cycle, Beta Oxidation, Acetogenesis(p) | B12 | Protein Degradation, Vitamin Production |
| H1N1.021 | Porticoccaceae | Piscivorous | Proteins | Glycolysis (p), Pyruvate Oxidation, Acetogenesis (p) | B9 | Protein Degradation, Vitamin Production |
| H1N1.022 | Alteromonadaceae | Piscivorous | Proteins, Fatty Acids, Cellulose, Hemicellulose, Starch | Glycolysis, Pyruvate Oxidation, TCA Cycle, Beta Oxidation, Acetogenesis (p) | B1, B2, B7, B9 | Protein Degradation, Vitamin Production |
| H1N1.025 | Alteromonadaceae | Piscivorous | Proteins, Starch | Glycolysis, Pyruvate Oxidation, Acetogenesis | B2, B9 | Protein Degradation, Vitamin Production |
| H1N1.026 | Sphingomonadaceae | Piscivorous | Proteins | Glycolysis, Pyruvate Oxidation, TCA Cycle (p), Acetogenesis (p) | B1, B7 | Protein Degradation, Vitamin Production |
| H1N1.028 | Maricaulaceae | Piscivorous | Proteins, Cellulose | Glycolysis (p), Pyruvate Oxidation, Acetogenesis (p) | B1, B7, B9 | Protein Degradation, Vitamin Production |
| H1N1.030 | Flammeovirgaceae | Piscivorous | Proteins, Fatty Acids, Cellulose, Hemicellulose, Starch, Chitin | Glycolysis, Pyruvate Oxidation, TCA Cycle (p), Beta Oxidation, Acetogenesis | B1, B7, B9 | Protein Degradation, Vitamin Production |
| H1N1.033 | GCA-2696645 | Piscivorous | Proteins, Cellulose | Glycolysis (p), Pyruvate Oxidation, TCA Cycle (p), Acetogenesis (p) | - | Protein Degradation |
| T4LTN.001 | Bacteroidaceae | Herbivorous | Cellulose, Hemicellulose, Starch, Proteins, Chitin | Glycolysis, Pyruvate Oxidation, TCA Cycle (p), H2-Producing Fermentation, Acetogenesis (p) | B1, B2, B7, B9 | Plant Matter Degradation, Vitamin Production |
| T4LTN.002 | Paludibacteraceae | Herbivorous | Cellulose, Proteins | Glycolysis, Pyruvate Oxidation, Acetogenesis (p) | - | Plant Matter Degradation |
| T4LTN.003 | Vibrionaceae | Herbivorous | Starch, Proteins, Fatty Acids, Chitin | Glycolysis, Pyruvate Oxidation, TCA Cycle (p), Nitrate reduction, Acetogenesis, Formate fermentation, Beta Oxidation | B1, B2, B7, B9 | Plant Matter Degradation, Vitamin Production |
| T4LTN.004 | Shewanellaceae | Herbivorous | Starch, Proteins, Fatty Acids | Glycolysis, Pyruvate Oxidation, TCA Cycle, Nitrate reduction, Acetogenesis, Formate fermentation, Beta Oxidation | B1, B2, B7, B9, B12 | Plant Matter Degradation, Vitamin Production |
| T4LTN.005 | Tannerellaceae | Herbivorous | Cellulose, Hemicellulose, Starch, Proteins, Chitin | Glycolysis, Pyruvate Oxidation, TCA Cycle (p), H<sub>2</sub>-Producing Fermentation, Acetogenesis (p) | B1, B2, B7, B9, B12 | Plant Matter Degradation, Vitamin Production |
| T4LTN.006 | Lachnospiraceae | Herbivorous | Cellulose, Hemicellulose, Starch, Proteins, Chitin | Glycolysis, Pyruvate Oxidation, Lactate fermentation, H<sub>2</sub>-Producing Fermentation, Acetogenesis (p) | B7, B12 | Plant Matter Degradation, Vitamin Production |
| T4LTN.007 | Alteromonadaceae | Herbivorous | Cellulose, Hemicellulose, Starch | Glycolysis, Pyruvate Oxidation, TCA Cycle (p), Nitrate reduction, Acetogenesis | B1, B2, B7, B9 | Plant Matter Degradation, Vitamin Production |
| T4LTN.009 | Alteromonadaceae | Herbivorous | Cellulose, Starch | Glycolysis, Pyruvate Oxidation, TCA Cycle (p), Acetogenesis | B1, B7 | Plant Matter Degradation, Vitamin Production |
| T4LTN.010 | Sphaerochaetaceae | Herbivorous | Proteins | Glycolysis, Pyruvate Oxidation, H<sub>2</sub>-Producing Fermentation, Acetogenesis (p) | - | - |
| T4LTN.011 | UBA1402 | Herbivorous | Hemicellulose | Glycolysis, Pyruvate Oxidation, TCA Cycle (p), Acetogenesis (p) | B9 | Plant Matter Degradation, Vitamin Production |
| T4LTN.012 | Vibrionaceae | Herbivorous | Starch, Proteins, Fatty Acids | Glycolysis, Pyruvate Oxidation, TCA Cycle, Nitrate reduction, Acetogenesis, Formate fermentation, Beta Oxidation | B1, B2, B7, B9, B12 | Plant Matter Degradation, Vitamin Production |
| T4LTN.014.2 | CAG-822 | Herbivorous | Starch | Glycolysis (p), Pyruvate Oxidation, Acetogenesis (p) | - | Plant Matter Degradation |
| T4LTN.014.3 | UBA6164 | Herbivorous | Simple Saccharides | - | - | - |
| T4LTN.015 | Ruminococcaceae | Herbivorous | Cellulose, Hemicellulose, Chitin | Glycolysis, Pyruvate Oxidation, H<sub>2</sub>-Producing Fermentation, Acetogenesis (p) | - | Plant Matter Degradation |
| **T4LTN.016** | Saccharospirillaceae | Herbivorous | Cellulose, Proteins, Fatty Acids | Pyruvate Oxidation, Nitrate reduction, Acetogenesis, Beta Oxidation | B1, B7, B9 | Plant Matter Degradation, Vitamin Production |
| **T4LTN.017** | Ruminococcaceae | Herbivorous | Simple Saccharides | Glycolysis, Pyruvate Oxidation, H<sub>2</sub>-Producing Fermentation, Acetogenesis (p) | B7 | Vitamin Production |
| **T4LTN.022** | Sphingomonadaceae | Herbivorous | Proteins | Glycolysis, Pyruvate Oxidation, TCA Cycle (p), Acetogenesis (p) | B1 | Vitamin Production |
| **T4LTN.023** | P3 | Herbivorous | Proteins | Glycolysis, Pyruvate Oxidation, H<sub>2</sub>-Producing Fermentation, Acetogenesis (p) | B1 | Vitamin Production |
| **Tr3LTN.002** | Shewanellaceae | Invertivorous | Proteins, Fatty Acids | Glycolysis, Pyruvate Oxidation, TCA Cycle, Nitrate reduction, Acetogenesis, Formate fermentation, Beta Oxidation | B1, B2, B7, B9, B12 | Protein Degradation, Vitamin Production |
| **Tr3LTN.005** | Rhodobacteraceae | Invertivorous | Proteins, Fatty Acids | Glycolysis, Pyruvate Oxidation, TCA Cycle (p), Beta Oxidation | B2, B9, B12 | Protein Degradation, Vitamin Production |
| **Tr3LTN.006** | Cellvibrionaceae | Invertivorous | Proteins, Fatty Acids | Glycolysis, Pyruvate Oxidation, Beta Oxidation | B1, B7, B9 | Protein Degradation, Vitamin Production |
| **Tr3LTN.008** | Arcobacteraceae | Invertivorous | Simple Saccharides | Glycolysis, Pyruvate Oxidation, TCA Cycle (p) | B1, B7 | Vitamin Production |
| **Tr3LTN.016** | Class: Bacilli | Invertivorous | Simple Saccharides | Glycolysis, Pyruvate Oxidation, H<sub>2</sub>-Producing Fermentation | - | - |
| **Tr3LTN.017** | UBA12223 | Invertivorous | Simple Saccharides | Glycolysis (p), Pyruvate Oxidation, TCA Cycle (p) | - | - |
| **Tr3LTN.026** | Bacteriovoracaceae | Invertivorous | Simple Saccharides | Glycolysis (p), Pyruvate Oxidation | - | - |

<sup>1</sup> Naming: H1N1, Hawfish; T4LTN, Yellow tang; Tr3LTN, Triggerfish
<sup>2</sup>Carbohydrate substrates are included if the MAG was capable of completely degrading it from start to finish. Starch is listed if the MAG possesses more than 5 amylase genes. Proteins are listed if the MAG possesses more than 10 exopeptidase genes. If the MAG could only produce β-glucosidase, simple saccharides is listed.
<sup>3</sup>For energy production, the pathway is included if the MAG can complete it entirely. If the MAG can complete an energy yielding/harvesting step of a pathway but not the entire process, the function is listed with a “p” (partial).

### Fatty Acids

All three hosts had MAGs that could carry out complete beta oxidation of fatty acids. The screen for fatty acid breakdown via beta oxidation included genes for acyl-CoA dehydrogenase, enoyl-CoA hydratase, 3-hydroxyacyl-CoA dehydrogenase, and acetyl-CoA acyltransferase. In the piscivorous gut community, 9/18 MAGs [003, 006, 007, 009, 012, 015, 018, 022, 030] were able to carry out fatty acid oxidation to completion to produce acetyl CoA (Figure 5a). These MAGs represent the *Gammaproteobacteria* (2), *Alphaproteobacteria* (3), and *Bacteroidia* (3) classes. In contrast, only 4/18 MAGs [003, 004, 012, and 016] from the herbivorous community and 3/7 MAGs [002, 005, and 006] from the invertivorous community could independently complete all of fatty acid oxidation (Figure 5b,c). All four MAGs from the herbivorous yellow tang microbiome that could complete beta oxidation were *Gammaproteobacteria*, and 3 of them were in the *Enterobacterales* order. These data highlight a potential enrichment of fatty acid oxidation capacities in the piscivorous hawkfish treatment, whose fish pellets contained high amounts of various fatty acids (Supplemental Table 1).

**Figure 5:**
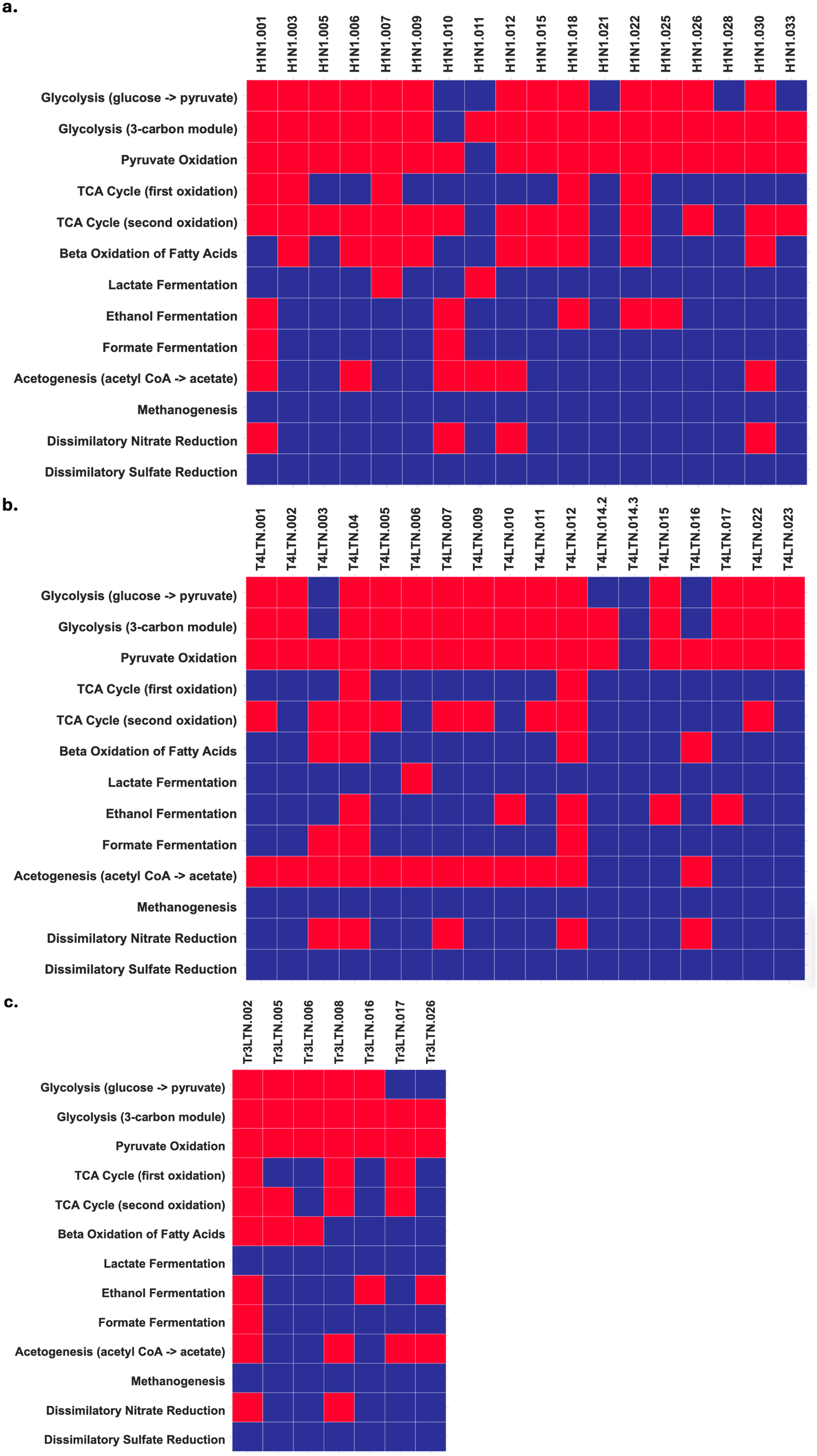
Heatmaps of metabolic functions that can be performed by recovered MAGs. Samples shown are a) piscivorous hawkfish, b) herbivorous yellow tang, and c) invertivorous triggerfish. Red indicates the ability to perform a metabolic pathway, while blue indicates inability.

### Energy Harvesting and Fermentation

MAGs recovered from all three samples were analyzed for their metabolic capacities. 13/18, 14/18 and 5/7 MAGs from the piscivorous, herbivorous, and invertivorous treatments, respectively, could carry all of glycolysis from glucose to pyruvate (Figure 5). Following glycolysis, 17/18, 17/18, and 7/7 MAGs from the piscivorous, herbivorous, and invertivorous gut communities could carry out pyruvate oxidation, bridging glycolysis to the citric acid cycle (Figure 5). 13/18, 6/18, and 4/7 MAGs from the piscivorous, herbivorous, and invertivorous guts, respectively, possessed more than 80% of the genes for the entire TCA cycle (Figure 5). Within each treatment, there are some MAGs that can complete individual energy-harvesting steps within the TCA cycle, but not the complete process, which may be a product of genome incompleteness (Table 3). Since the piscivorous hawkfish sample contained a higher proportion of MAGs that could carry out the TCA cycle, particularly the second oxidation reactions, these data could suggest that these MAGs are more predominantly metabolizing the amino acids from their proteinaceous diet via central carbon metabolism.

The central carbon pathways observed in these MAGs produce reduced electron carriers that need to be oxidized in order to sustain the catabolic pathways. From the piscivorous treatment, 4/18 MAGs [001, 010, 012, and 030] could perform dissimilatory nitrate reduction to ammonia through a nitrite intermediate as a mechanism for anaerobic respiration (Figure 5a). 5/18 MAGs [003, 004, 007, 012, and 016] from the herbivorous treatment are dissimilatory nitrate respirers, while 2/7 MAGs [002 and 008] from the invertivorous treatment can carry out nitrate respiration (Figure 5b-c). Lactate fermentation was rare among all three treatments, with only 2/18, 1/18, and 0/7 MAGs from the piscivorous, herbivorous, and invertivorous diets, respectively, being able to produce a L-lactate dehydrogenase (Figure 5).

All three gut communities demonstrated an ability to perform the latter half of the acetogenesis pathway to reduce acetyl-CoA to acetate. From the piscivorous treatment, 6/18 MAGs had genes for both phosphate acetyltransferase and acetate kinase, synthesizing ATP Figure 5a; Supplemental Figure 2). Similarly, 11/18 MAGs from the herbivorous diet and 4/7 MAGs from the invertivorous diet could produce these enzymes (Figure 5b-c; Supplemental Figures 3-4). Using the acetate products, 5/18, 7/18, and 3/7 MAGs from the piscivorous, herbivorous, and invertivorous communities, respectively, could produce alcohol dehydrogenase homologs for the production of ethanol and consumption of reduced NADH (Figure 5; Supplemental Figures 2-4).

Alternatively, all three treatments had MAGs that had genes for formate acetyltransferase and formate dehydrogenase, which are involved in the fermentation of formate to carbon dioxide and hydrogen gas. From the piscivorous treatment, 3/18 MAGs could produce formate acetyltransferase while 8/18 MAGs could produce formate dehydrogenase (Figure 5a; Supplemental Figure 2). In contrast, 11/18 MAGs from the herbivorous diet could produce formate acetyltransferase, while only 3/18 MAGs could produce formate dehydrogenase (Figure 5b; Supplemental Figure 3). From the invertivorous diet, 4/7 MAGs possessed formate acetyltransferase genes, while 3/7 MAGs possessed genes for formate dehydrogenase (Figure 5c; Supplemental Figure 4). None of the MAGs in this study could perform dissimilatory sulfate respiration or methanogenesis.

### Vitamin and Hydrogenase Production

All communities had MAGs that could produce vitamins B1, B2, B7, B9, and B12. Thiamine (B1) could be produced by 7/18, 10/18, and 3/7 MAGs from the piscivorous, herbivorous, and invertivorous communities, respectively (Table 3). Riboflavin (B2) could be synthesized by 4/18, 6/18, and 2/7 MAGs from the piscivorous, herbivorous, and invertivorous hosts guts, respectively (Table 3). Biotin (B7) could be synthesized by 12/18, 10/18, and 3/7 MAGs from the piscivorous, herbivorous, and invertivorous diet communities, respectively, using at least one of the various biotin biosynthetic pathways (Table 3). Hydrotetrafolate, a precursor to folic acid (B9), could be produced by 14/18, 9/18, and 3/7 MAGs from the piscivorous, herbivorous, and invertivorous hosts guts, respectively (Table 3). Cobalamin (B12) could be made by 7/18, 4/18, and 2/7 MAGs from the piscivorous, herbivorous, and invertivorous hosts guts, respectively (Table 3).

From the piscivorous gut community, the only MAG capable of producing a component of a nickel-iron hydrogenase is 001 (*Photobacterium aphoticum*), being able to produce a periplasmic Group 1 hydrogenase, which is generally responsible for catalyzing steps in anaerobic respiration using metals, nitrates, or sulfates as terminal electron acceptors (Supplemental Figure 5). Similarly only 1 MAG [016] from the invertivorous treatment produces a group A1 and A3 iron-iron hydrogenase, which produces fermentative hydrogen gas (Supplemental Figure 5). Also from the invertivorous diet, 2/7 MAGs [002 and 008] from this diet produce group 1 nickel-iron hydrogenases., which also act as respiratory electron receptors (Supplemental Figure 5).

In contrast, MAGs from the herbivorous treatment synthesize various iron and nickel hydrogenases. 6/18 MAGs [005, 010, 015, 017, 023, and 006] produce group A1 and A3 iron-iron hydrogenases (Supplemental Figure 5). 7/18 MAGs [001, 005, 010, 015, 017, 023, and 006] produce group B iron-iron hydrogenases involved in hydrogen gas fermentation (Supplemental Figure 5). 4/18 MAGs [010, 015, 017, and 006] produce group C1 iron-iron hydrogenases (Supplemental Figure 5). Only 2/18 MAGs [017 and 006] produce group C2 iron-iron hydrogenases, which are transcriptional regulators (Supplemental Figure 4). 4/18 MAGs [010, 015, 023, and 006] produce group C3 iron-iron hydrogenases (Supplemental Figure 5). 3/18 MAGs [001, 004, and 005] produce group 1 nickel-iron hydrogenases, also transcriptional regulators (Supplemental Figure 5). Within the herbivorous yellow tang gut, MAGs 006, 010, 015, and 017 produce the greatest variety of hydrogenases, and 3 of them belong to the *Clostridia* class within *Firmicutes*.

## Discussion

The present study used metagenome-assembled genomes recovered from the feces of tropical reef fishes that were collected from the wild and housed in aquaria under controlled diets. We found evidence of core functions in the examined microbiomes. Our study benefited from the aquarium-based approach, as we had defined food sources that the fish were fed for weeks in captivity, compared to the uncontrolled diet that may be found in nature. Core functions include protein degradation, fatty acid oxidation, central carbon metabolism, nitrate respiration, acetogenesis, and formate oxidation. Considering our sampling design and dataset limits, additional functions may be possible in fish guts and more taxa should be closely examined; however, our results show that previous work based on small subunit ribosomal sequences or the interpretation of metabolism based solely on taxonomic data is a limited view of a fish gut microbiome.

Despite the number of shared functions, diet types also showed selective trends. The microbial MAGs associated with the piscivorous and invertivorous diets were the most functionally similar, as expected, considering the compositions of the fish pellets and shrimp diets, consistent with the model of diet-based microbiomes (34). Both diets were high in protein and fat content, and the MAGs recovered from these diets showed high gene copy numbers for exopeptidases and the abilities to carry out beta oxidation of fatty acids. The piscivorous gut community, although specialized for the degradation of proteins and fatty acids, could also degrade plant matter. It could be that plant material is indirectly ingested during fish consumption. In contrast, the invertivorous triggerfish is primarily digesting shrimp, which likely do not contain substantial amounts of complex plant polysaccharides in their intestines. Therefore, the triggerfish MAGs encode for digesting proteins and fats with some versatility for simple sugars. As a caveat, examination of additional MAGs after additional sequencing effort could show a broader digestion ability for the triggerfish.

As expected, the herbivorous yellow tang gut MAGs demonstrated a strong ability to degrade complex plant polysaccharides, likely due to their nori seaweed diet. Unexpectedly, the MAGs recovered from the invertivorous triggerfish diet demonstrated an inability to degrade chitinous compounds. This contrasts with previous results which found that rainbow trout being fed chitinaceous insect exuviae had gut microbiomes enriched in chitinolytic bacterial genera; however, these results were based on predictive metagenomes based solely on 16S rRNA gene amplicons (35). Exochitinase production has been observed in various fish species being fed insect diets, although the source of these enzymes (host versus gut microbiome) was not specified (36, 37).

With the higher fatty acid diets of the piscivorous hawkfish and the invertivorous triggerfish, their MAGs carried a strong signature of beta oxidation. The beta oxidation of fatty acids is a reductive cycle that rapidly degrades long chain fatty acids to directly produce ATP and acetyl-CoA. This high energy-yielding process explains its abundance in MAGs recovered from these treatments, while the herbivorous yellow tang gut microbial community had fewer MAGs capable of fatty acid degradation. Instead of relying on this high yield process, our data suggest that the herbivorous gut community relies more on mixed acid fermentation and acetate-producing pathways for the production of ATP and recycling of reduced electron carriers. This establishes a major difference in the proteinaceous and fatty acid diet economy expected in the piscivorous and invertivorous treatments and the polysaccharide diet economy expected in the herbivorous treatment.

Further supporting the polysaccharide economy expected in the herbivorous yellow tang gut, the MAGs collected from this treatment possessed a deep redundancy and breadth of hydrogenases. These hydrogenases varied in function, including being final electron acceptors in anaerobic respiration and hydrogen gas production at the end of fermentation. Since the herbivorous treatment is associated with higher concentrations of fibrous substrates, the constituent sugars would rely on electron mediated processes to degrade the substrates and recycle cofactors. The higher proportion of MAGs that produce hydrogenases supports this expectation, with these hydrogenases playing a putative role in balancing electron budgets and gradients to support respiration and fermentation pathways.

A limitation of this study that is worth noting is the possibility for there to be host-specific factors such as gut physiology and chemistry that could also be selecting for taxon and function-specific MAGs. While the present study found substantial evidence for the linkage between diet inputs and community composition, it has been shown that host species may harbor distinct microbial communities (7). Future research should study model fish species that have adaptable diets or whose diets change with development (38). Then, diets could be shifted and changes in the microbial community composition could be followed as a function of time and treatment. The methodology used here shows that fish under treatment can have their microbiome analyzed while remaining viable for longer treatments of study. If these diet changes are considered in the context of ecological disturbances, these studies would be indicative of the likelihood of adjusting to rapidly changing environments, which is important to understand given the cascade of environmental changes being mediated by anthropogenic activity.

We recognize that keeping fish viable for this study meant that internal shifts of microbes along gut structures were unable to be observed (5, 39, 40). As such, these data serve as a preliminary basis for further studies whose experimental design could involve more intensive sampling. However, our data does show strongly that single-gene taxonomic analyses fall short of determining differences in fish gut microbiomes, and many lineages may be more or less represented based on the method of molecular analysis. The ability to sample the full metagenome and build MAG-based models of metabolism shows that fish guts are more similar in functional potential than expected. In order to get insight into the functions of these MAGs within these microbiomes, a full study should be done that addresses biomass abundance within the gut, potentially by transcriptomics or proteomics, which can provide knowledge of how these microbes are behaving and regulating the expression of their genomes.

## Materials and Methods

### Fish Rearing, Diets and Sample Collection

The yellow tangs, triggerfish, and hawkfish used for this study were wild caught and obtained from Live Aquaria ® (Rhinelander, WI) and cared for according to IACUC approval through the University of Delaware (A1319). Fish were held in 37L aquaria on a large recirculating system at the UD Lewes Campus (Lewes, DE). Hawkfish (n=4) were kept individually in aquarium tanks due to aggressive behavior. Yellow tangs (n=5) and triggerfish (n=5) were kept in communal aquarium tanks with their same species. Hawkfish are piscivorous fish and were fed commercial fish meal pellets (Sustainable Aquatics Dry Hatchery Diet, Jefferson City TN). Yellow tangs are herbivorous fish and were fed commercially purchased nori seaweed (sushi wrapping, various vendors). Triggerfish are invertivores and were fed commercially purchased mysis shrimp (San Francisco Bay Brand Frozen Mysis Shrimp). We opted to collect gut microbiomes via fecal collection to preserve the fish for other experiments (41).

When collecting feces, which occurred multiple times due to the fish not being sacrificed, fish were placed in individual 2.5L aquaria. To minimize a seawater microbial signal, artificial seawater was created with Milliq-filtered water and Instant Ocean (Instant Ocean; Blacksburg, VA) to 33 ppm salinity. Fish were moved to these tanks 10 min after feeding and remained there overnight with air stones and no shared water flow to ensure that water quality remained high. After 12-15 hours, the fish were placed back in their 37L tanks. From each small tank, 1500 mL of water was sampled. The tank waters were filtered through an 8.0 micron cellulose filter (Merck Millipore; Burlington, MA). This filter size was selected to ensure fecal particles were preferentially collected instead of free-living bacteria in the water. The filters were labeled and placed in a 60mm x 15mm petri plate with a stackable lid and stored in a -80C freezer until used for DNA extractions.

### Chemical Analysis of Food Sources

To determine major dietary differences between the fish-meal pellets and mysis shrimp, 20g of crushed, freeze dried mysis shrimp and 20g of fish-meal pellets were shipped to NP Analytics Laboratory (St. Louis, MO) for testing. Nutrition information for nori seaweed was retrieved from the commercial packaging.

### DNA Extraction, Amplicon and Metagenomic Sequencing

DNA was extracted from the feces samples collected on the filter using the DNeasy PowerSoil Kit (Qiagen, Valencia CA). A quarter of the filter paper was used for the extractions. After each extraction, the amount of DNA in each sample was measured using the Qubit dsDNA HS assay kit with the Qubit fluorometer (Invitrogen; Waltham, MA). The DNA was stored in a - 20C freezer. Samples with the highest DNA yield were selected from each species and sent to the University of Connecticut MARS facility for amplification and sequencing of the 16S rRNA gene (primers 515F, 806R) (42) using the Illumina MiSeq platform via their standard protocol. After 1 month of daily feeding, additional samples were taken in the same manner, processed the same way and sent to the University of Delaware DNA Sequencing and Genotyping Center at the Delaware Biotechnology Institute (Newark, DE) for library creation and sequencing on an Illumina HiSeq 2500.

### Data Accessibility Statement

Raw metagenome data associated with these samples have been deposited in the NCBI database under BioProject 1112800. Amplicon sequences are under BioProject PRJNA1174778. Individual metagenome assembled genomes (MAGs) and annotation files are available at https://figshare.com/projects/FishGutMAGs_AEM2024/223746.

### Amplicon Sequence Quality Control and Processing

Raw forward and reverse amplicon sequences were paired and processed using 16S rRNA analysis pipeline in MOTHUR version 1.46.1 (43). Paired sequences were trimmed, ambiguous nucleotides were removed, and then operational taxonomic units (OTUs) with a 3% dissimilarity were created. OTUs were then aligned and classified using the Silva138 database version 138 (44).

### Metagenome Quality Control, Assembly and Binning

Data processing, including reads quality checking and trimming, was performed using Nesoni (45) under default parameters and applying a quality score of Q20. Qualified reads for each dataset were assembled into contigs using IDBA-UD (46), metaSPAdes (47), and Megahit v.1.1 (48) with default settings and k-mer lengths varying from 27 to 117. Community composition was determined via annotation and retrieval of all 30S ribosomal protein (rps3) sequences from assembled contigs. Reads were mapped to each rps3 gene using gbtools (49) and BLAST (50) against UniProt (51) to identify the taxonomy of the gene. Community composition was determined by the coverage of each gene compared to the total read coverage of all rps3 genes in the dataset. Contigs longer than 1000 bp were binned using binning tools MaxBin2.0 (52), Metabat (53), and Metabat2 (54). The resulting metagenome-assembled genomes (MAGs) were quality assessed using CheckM (55) with the “lineage_wf” option. Only bacterial MAGs, meeting criteria of ≥ 50% completion and ≤ 20% contamination, were retained. These MAGs were further refined by removing outlier contigs based on t-SNE signatures, GC content, taxonomic assignments, and differential coverages. Coverage calculations were performed by mapping trimmed reads against the contigs using BBMap v.37.61 (56). MAGs with < 10% contamination were deemed acceptable, while those with 10-20% contamination were manually cleaned again.

### Phylogenetic Placement

The phylogenetic placements of the MAGs were estimated by creating a phylogenomic tree from concatenated ribosomal proteins, aligned using Muscle v3.8 (57) and analyzed with FastTree v2.1 (58). MAG identities were confirmed by calculating average nucleotide identity (ANI) and average amino acid identity (AAI) with custom reference genomes closely related to the screened lineages based on their single marker gene(s) closest matches. Taxonomic profiles were verified using GTDB-Tk v2.1.0 (59).

### Metabolic Reconstruction

Protein sequences from all MAGs were predicted using Prodigal v2.6.3 (60) using the default translation table 11. The collective metabolic pathways for all MAGs present in each depth were deduced via screening the predicted proteins against a custom hidden Markov model database composed of 126 key metabolic genes covering most of the substrate utilization, biosynthesis and energy metabolism related pathways (61). These pathways were further assessed for completion through querying against the Kyoto Encyclopedia of Genes and Genomes (KEGG) (https://www.genome.jp/kegg/) (62) database using the BlastKoala tool (63). The pathway was deemed complete if the MAG encodes for at least 80% of the enzymes and other involved proteins of a given pathway. We validated our results using METABOLIC-C (64). METABOLIC-C integrates gene calling and annotation based on HMM-based databases, motif validation, and annotation via specific databases.

### Substrate Degradation Analysis

CAZyme sequences were identified via HMM searches against dbCAN V8 (65). Enzymes involved in the degradation of cellulose, hemicelluloses (xylan, mannan, and xyloglucan), and starch were identified as presented in Kumla et al. (66). Enzymes for the degradation of chitin were identified as presented in De Tender et al. (67). Corresponding CAZyme families for these enzymes were identified on the Carbohydrate-Active enZYmes Database (http://www.cazy.org/) (68). Exopeptidases were identified using the MEROPS Peptidase Database (https://www.ebi.ac.uk/merops/) (69). Enzymes involved in lactate, acetate, and ethanol fermentation pathways were organized as described in the EcoCyc Database (https://ecocyc.org/) (70). Formate fermentation enzymes were identified as described in Knappe & Sawers (71). Hydrogenase enzyme sequences were identified through specific HMM searches against the Kofam database V2024-01-01 (72). Hydrogenase functions were identified on the HydDB database (https://services.birc.au.dk/hyddb/) (73).

### Statistical Analysis

To visualize clustering patterns in substrate degradation enzymes in response to host diets, a principal component analysis of all screened substrate pathways was performed on all samples using the prcomp function in R Version 4.3.2 (74). The total number of gene copies for all enzymes involved in the degradation of proteins, cellulose, hemicellulose, starch, and chitin normalized to genome completeness and a binary system for the presence/absence of fatty acid degradation were scaled and used for PCA processing. The loadings of the first two components are reported, explaining 77.0% of detected variance. PCA data were visualized using the fviz_pca_biplot function from the factoextra package (75). To test if the centroids of diet treatments were different, a permutational multivariate analysis of variance (PERMANOVA) was performed using the adonis2 function from the vegan package (76). To assess the homogeneity of multivariate dispersions across diets, the betadisper and permutest functions were used to compute the distances from group centroids to individual samples with 999 permutations.

To determine if MAGs recovered from each treatment differed in the number of substrate degrading gene copies, the total gene copies for each degradation pathway normalized to MAG genome completeness were totaled. Then, a Kruskal Wallis test was performed to determine if there were significant differences among the three diets using the kruskal.test function. If significant, a post-hoc Wilcoxen test was performed to identify significantly different pairwise comparisons between diets.

### Ethics Statement

The handling and care protocols of the fish used in this study were approved by the University of Delaware Institutional Animal Care and Use Committee (A1319), AUP #1319-2019-2.

## Acknowledgements

We would like to thank our colleagues at UD in Lewes for assistance with fish care and sampling. This work was supported by The Mindlin Foundation, funding to CR via the UD Graduate Scholars Fellowship, funding to KK and JFB from the WM Keck Foundation and funding to IF and JFB by ExxonMobil Research and Engineering Company. DW was supported as a UD School of Marine Science & Policy scholar. Support from the University of Delaware CBCB Bioinformatics Data Science Core Facility (RRID:SCR_017696), including use of the BIOMIX and BioStore computational resources was made possible through funding from Delaware INBRE (NIGMS P20GM103446), NIH Shared Instrumentation Grant (S10OD028725) the State of Delaware, and the Delaware Biotechnology Institute.

