## Supplemental Tables and Figures for "Comparative metagenomics of tropical reef fishes show conserved core gut functions across hosts and diets with diet-related functional gene enrichments"

**Supplemental Table 1**: Food analysis between mysis shrimp and fish-meal pellets from NP Analytics Laboratory analyzing protein by combustion, fatty acids by FANL/FANO, fiber, volatile organic acids, and minerals.

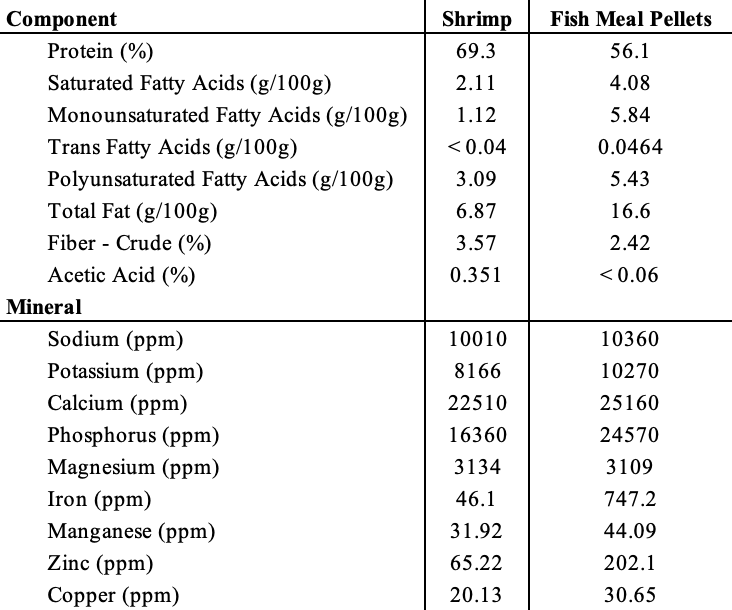

**Supplemental Table 2**: Food compositional analysis between mysis shrimp and fish-meal pellets from NP Analytics Laboratory analyzing fatty acids by FANL/FANO.

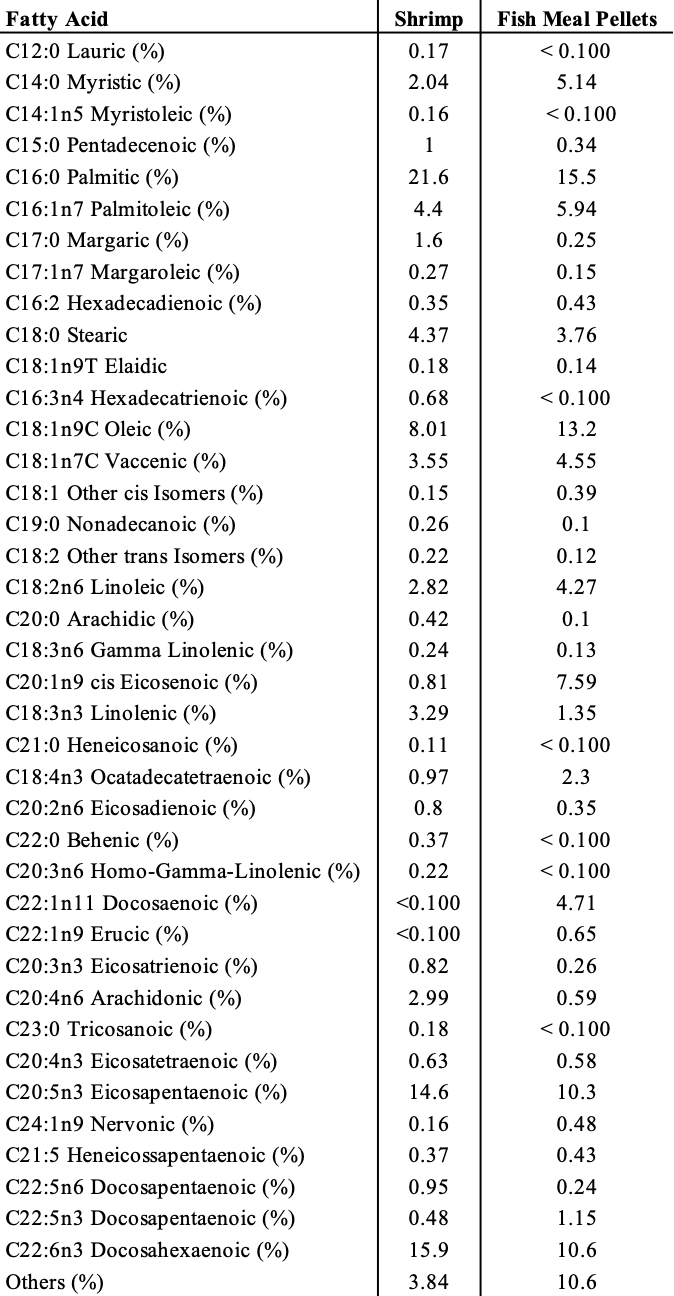

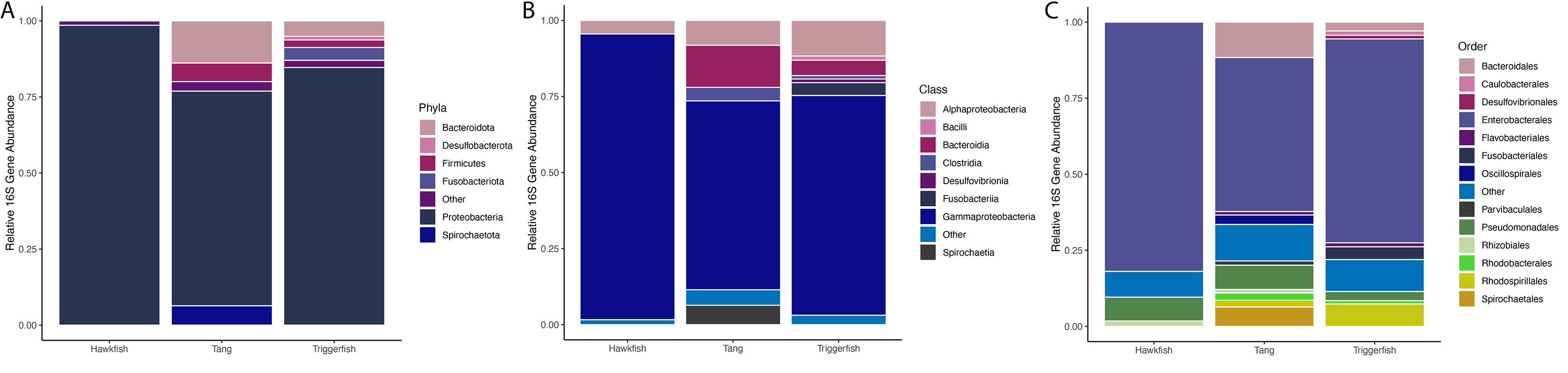

**Supplemental Figure 1:** Relative abundance of 16S rRNA genes in each fish listed at the A) Phylum B) Class and C) Order levels.

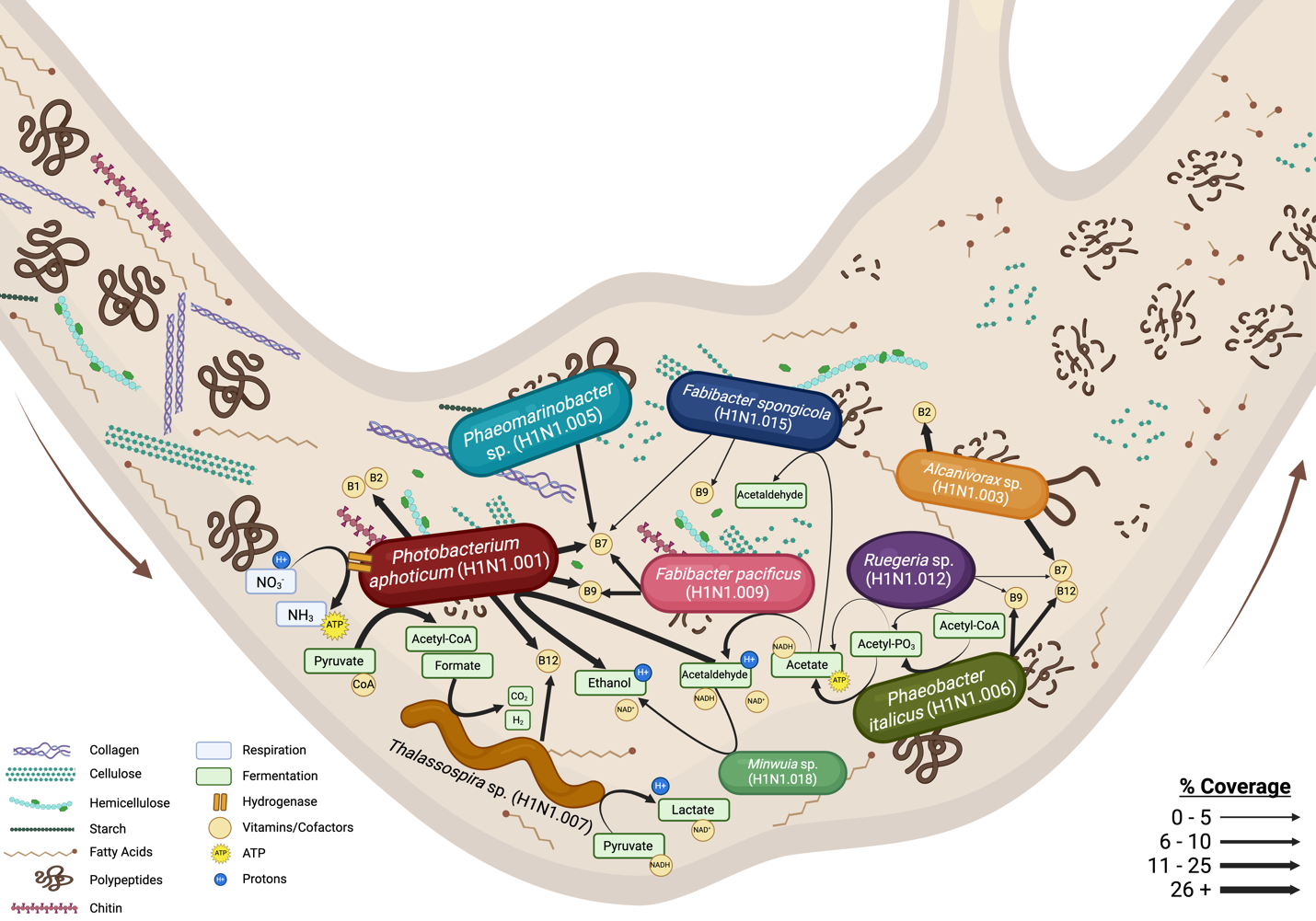

**Supplemental Figure 2:** Diagrammed annotations within the picivorous hawkfish MAGs. Arrows are sized to represent the MAG coverage.

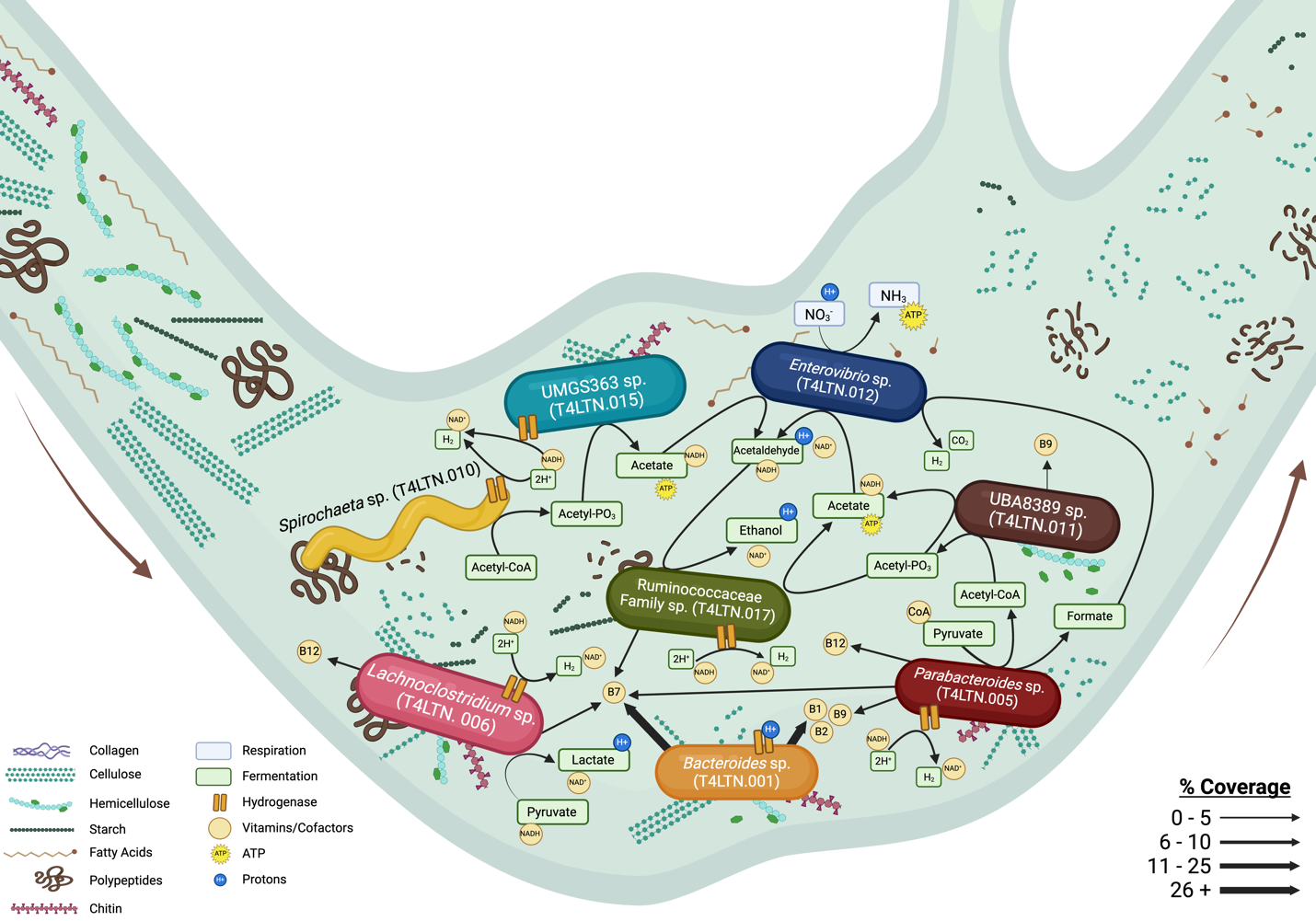

**Supplemental Figure 3:** Diagrammed annotations within the herbivorous yellow tang MAGs. Arrows are sized to represent the MAG coverage.

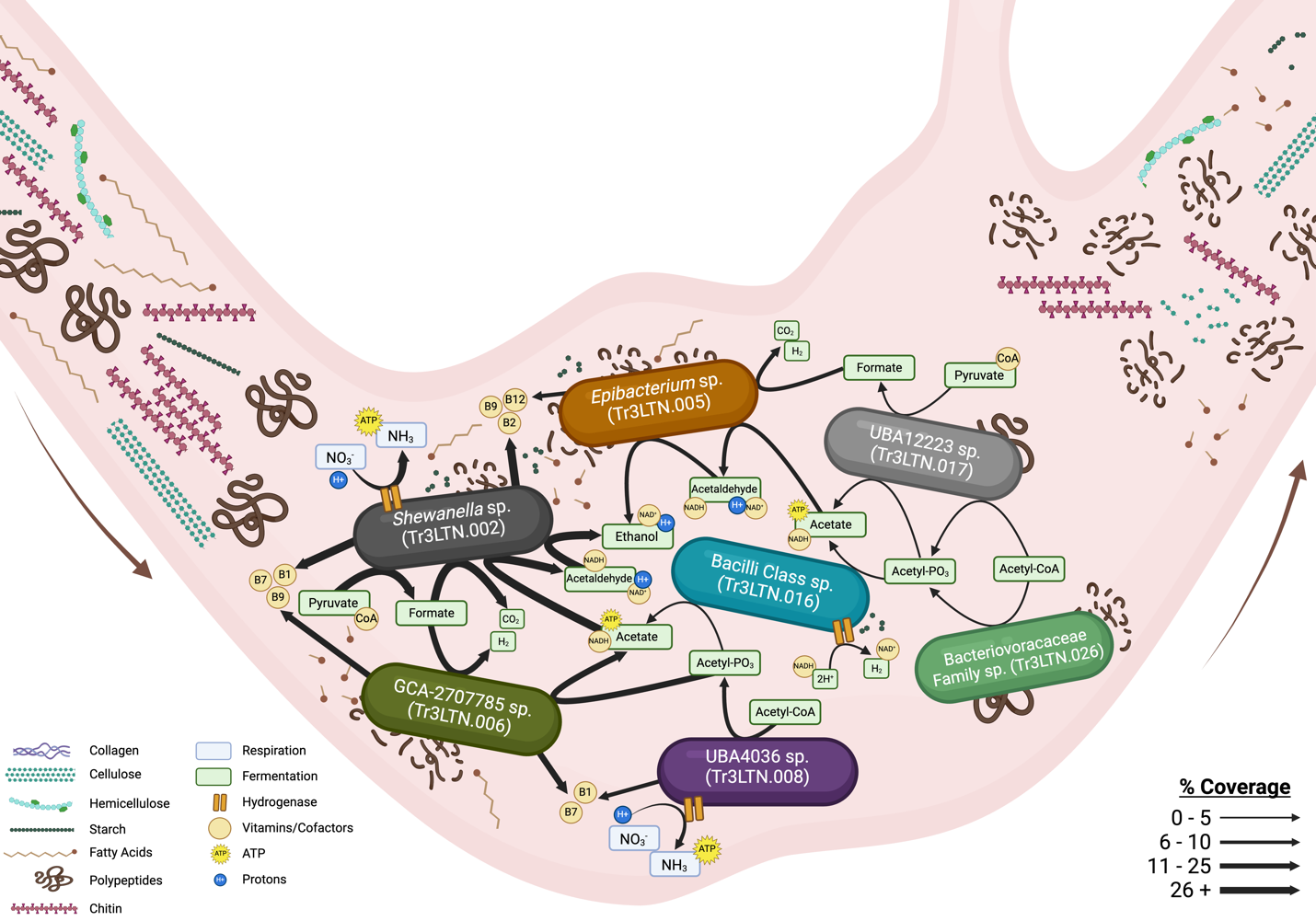

**Supplemental Figure 4:** Diagrammed annotations within the invertivorous triggerfish MAGs. Arrows are sized to represent the MAG coverage.

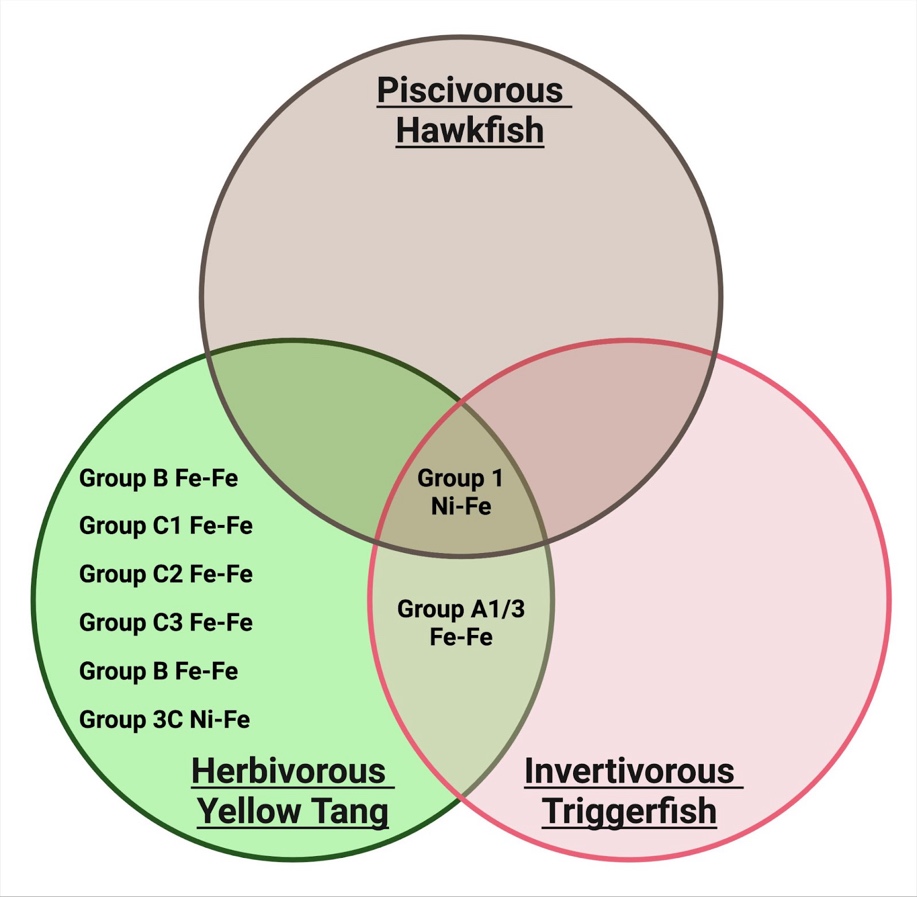

**Supplemental Figure 5**: Hydrogenase groups encoded in MAGs from each gut microbiome.
